# TopBP1 assembles nuclear condensates to switch on ATR signalling

**DOI:** 10.1101/2020.10.05.326181

**Authors:** Camilla Frattini, Alexy Promonet, Emile Alghoul, Sophie Vidal-Eychenie, Marie Lamarque, Marie-Pierre Blanchard, Serge Urbach, Jihane Basbous, Angelos Constantinou

## Abstract

ATR checkpoint signalling is crucial for cellular responses to DNA replication impediments. Using an optogenetic platform, we show that TopBP1, the main activator of ATR, self-assembles extensively to yield micron-sized condensates. These opto-TopBP1 condensates are functional entities organized in tightly packed clusters of spherical nano-particles. TopBP1 condensates are reversible, occasionally fuse and co-localise with TopBP1 partner proteins. We provide evidence that TopBP1 condensation is a molecular switch that amplifies ATR activity to phosphorylate checkpoint kinase 1 (Chk1) and slowdown replication forks. Single amino acid substitutions of key residues in the intrinsically disordered ATR-activation domain disrupt TopBP1 condensation and, consequently, ATR/Chk1 signalling. In physiologic salt concentration and pH, purified TopBP1 undergoes liquid-liquid phase separation *in vitro*. We propose that the actuation mechanism of ATR signalling is the assembly of TopBP1 condensates driven by highly regulated multivalent and cooperative interactions.

## Introduction

The primary structure of DNA is subjected to constant chemical alterations caused by spontaneous decay, endogenous metabolites and environmental genotoxic agents (Friedberg et al., 2006; Lindahl, 1993), hence organisms have evolved multiple DNA repair mechanisms to ensure genome integrity and survival (Ciccia and Elledge, 2010; Jackson and Bartek, 2009; Tubbs and Nussenzweig, 2017). DNA damage sensors, protein scaffolds and DNA processing activities accumulate at DNA damage sites to form spatially defined and reversible structures not delimited by membranes commonly referred to as nuclear foci (Garcia-Higuera et al., 2001; Lisby et al., 2001; Maser et al., 1997; Park et al., 1996). Much still remains to be understood about the molecular forces that drive the formation of DNA damage foci and their functional consequences in the DNA damage response.

In recent years, application of the principles of polymer chemistry to biological molecules has accelerated spectacularly our understanding of how membraneless compartments assemble and regulate their compositions and functions (Banani et al., 2017; Bracha et al., 2019; Hyman and Simons, 2012; Shin and Brangwynne, 2017; Soding et al., 2020). These self-organized micron-scale structures, called biomolecular condensates, assemble via multiple weak, cooperative and dynamic interactions and compartmentalize proteins and nucleic acids. The self-organization of multivalent molecules can yield a rich repertoire of higher-order structures with diverse physical properties, variable size, and no defined stoichiometry of constituent proteins. Whereas nucleic acids serve a seeding platform for the self-organization of soluble proteins (Mao et al., 2011; McSwiggen et al., 2019), soluble bridging factors can cross-link chromatin segments to compartmentalize chromatin via a process of polymer-polymer phase separation (Erdel et al., 2020). Moreover, increasing evidence indicates that diverse multivalent protein scaffolds self-organize via liquid-liquid phase separation, a process of de-mixing that yields a condensed phase enriched in the protein and a dilute phase (Banani et al., 2017; Shin and Brangwynne, 2017). Biomolecular condensates are functional hubs implicated in diverse cellular processes, including innate immune signalling (Du and Chen, 2018), microtubule nucleation (Woodruff et al., 2017), transcription (Boija et al., 2018; Kwon et al., 2013; Lu et al., 2018; Sabari et al., 2018) and adaptative stress responses (Franzmann and Alberti, 2019; Franzmann et al., 2018; Riback et al., 2017). Recent evidence indicates that protein phase separation can also occur at DNA damage sites. Upon laser micro-irradiation, the activity of poly(ADP-ribose) polymerase 1 seeds the condensation of the prototypical liquid-liquid phase separation protein FUS at damaged chromatin (Altmeyer et al., 2015; Patel et al., 2015). The DNA repair protein 53BP1 phase separates at double-strand DNA breaks (Kilic et al., 2019; Pessina et al., 2019), and the condensation of 53BP1 promotes induction of p53 and p21 (Kilic et al., 2019).

To explore the mechanisms and functional consequences of biomolecular condensates in the DNA damage response (DDR), we studied Topoisomerase IIβ-binding protein (TopBP1), an essential factor in the DDR pathway and a prototype of protein scaffolds composed of multiple modular interaction domains. TopBP1 features nine repetitions of a well-folded protein-protein interaction motif, the BRCA1 C terminus domain (BRCT), and an ATR activation domain (AAD), located between BRCT6 and BRCT7, which is intrinsically disordered. In a highly regulated manner, TopBP1 brings together different sets of proteins to form distinct protein complexes involved in DNA replication initiation (Hashimoto and Takisawa, 2003; Makiniemi et al., 2001), DNA replication stress signalling (Kumagai et al., 2006; Mordes et al., 2008), DNA repair (Broderick et al., 2015; Leimbacher et al., 2019; Liu et al., 2017; Moudry et al., 2016) and transcription regulation (Liu et al., 2009; Wright et al., 2006). In S phase, TopBP1 is the main activator of the master checkpoint kinase ATR (Kumagai et al., 2006; Mordes et al., 2008). ATR and its effector kinase Chk1 are crucial for cellular responses to DNA damage and DNA replication impediments (Ciccia and Elledge, 2010; Marechal and Zou, 2013; Saldivar et al., 2017). ATR/Chk1 signalling ensures cell and organismal survival through coordination of DNA repair and DNA replication with physiological processes, including cell cycle progression and transcription (Saldivar et al., 2017). Studies using Xenopus egg extracts have largely contributed to defining the orchestrated set of events leading to ATR activation (Acevedo et al., 2016; Byun et al., 2005; Duursma et al., 2013; Kumagai et al., 2006; Van et al., 2010). ATR is recruited to DNA lesions and replication intermediates via the ATR interacting protein ATRIP, which binds RPA-covered single-stranded DNA (Zou and Elledge, 2003). TopBP1 interacts with ATRIP, with ATR and with the RAD9-RAD1-HUS1 (9-1-1) clamp and activates ATR (Delacroix et al., 2007; Duursma et al., 2013; Kumagai et al., 2006; Mordes et al., 2008; Yan and Michael, 2009). Here, we used the conceptual framework born from studies of biomolecular condensates to gain fresh insights into the activation mechanism of the ATR/Chk1 signalling pathway. Using a combination of optogenetic and biochemical approaches, we demonstrate that TopBP1 can self-assemble extensively to yield functional biomolecular condensates. We show that purified TopBP1 has an intrinsic capacity to undergo liquid phase separation *in vitro*. Moreover, we provide evidence that TopBP1 self-organization into micron-sized compartments in living cells activates ATR/Chk1 signalling and slows down the progression of replication forks. Our data indicate that essential responses to DNA replication impediments emerge from TopBP1-driven assembly of nuclear condensates.

## Results

### TopBP1 drives the formation of micron-sized nuclear condensates

To probe the capacity of TopBP1 to self-organize into biomolecular condensates, we fused TopBP1 to cryptochrome 2 (Cry2) of *Arabidopsis thaliana*, a protein that oligomerises upon exposure to 488nm light (Bugaj et al., 2013) (Fig. 1A). This optogenetic system allows to control the nucleation of biomolecular condensates in space and time (Bracha et al., 2018; Shin et al., 2017; Shin et al., 2018), and evaluate the functional consequences of protein condensation (Kilic et al., 2019; Sabari et al., 2018). When fused to a protein scaffold that can undergo phase separation, the light-responsive Cry2 protein seeds the formation of biomolecular condensates that are held together by multivalent, weak and cooperative interactions (Kilic et al., 2019; Shin et al., 2017; Zhang et al., 2019). We induced the expression of TopBP1 fused to mCherry and Cry2 (named opto-TopBP1) with doxycycline in Flp-In HEK293 cells. Overall, the expression level of recombinant opto-TopBP1 was similar to the level of endogenous TopBP1 (Suppl. Figure 1A). At this level of expression, opto-TopBP1 remained in a diffuse state and was not directly detectable by fluorescence microscopy (Figure 1B, light OFF panel). Upon exposure of these cells to an array of blue-light LEDs during 3 minutes of light-dark cycles (4s light followed by 10s dark), we observed multiple and distinct opto-TopBP1 foci in the nuclei, specifically (Figure 1B, light ON panel). By contrast, we did not detect any foci in blue-light exposed cells expressing the opto-module (mCherry-Cry2) alone, nor in cells expressing the checkpoint clamp subunit opto-RAD9 (RAD9-mCherry-Cry2) (Suppl. Figure 1B). We compared cells expressing WT or W1145R opto-TopBP1. This mutation locates inside TopBP1 ATR activation domain (AAD), an intrinsically disordered domain required to directly stimulate the kinase activity of ATR (Kumagai et al., 2006). The tryptophan to arginine substitution W1138R in the ATR activation domain of *Xenopus Laevis* TopBP1 suppresses its capacity to activate *Xenopus* ATR (Kumagai et al., 2006). Furthermore, the homologous substitution W1147R is embryonic lethal in mice (Zhou et al., 2013). Upon optogenetic activation, W1145R TopBP1 expressing cells exhibited a markedly reduced number of TopBP1 condensates in comparison with wild-type TopBP1 (Figure 1B), suggesting that this aromatic residue plays an important role in TopBP1 higher-order assembly or the growth of TopBP1 condensates.

**Figure 1.**
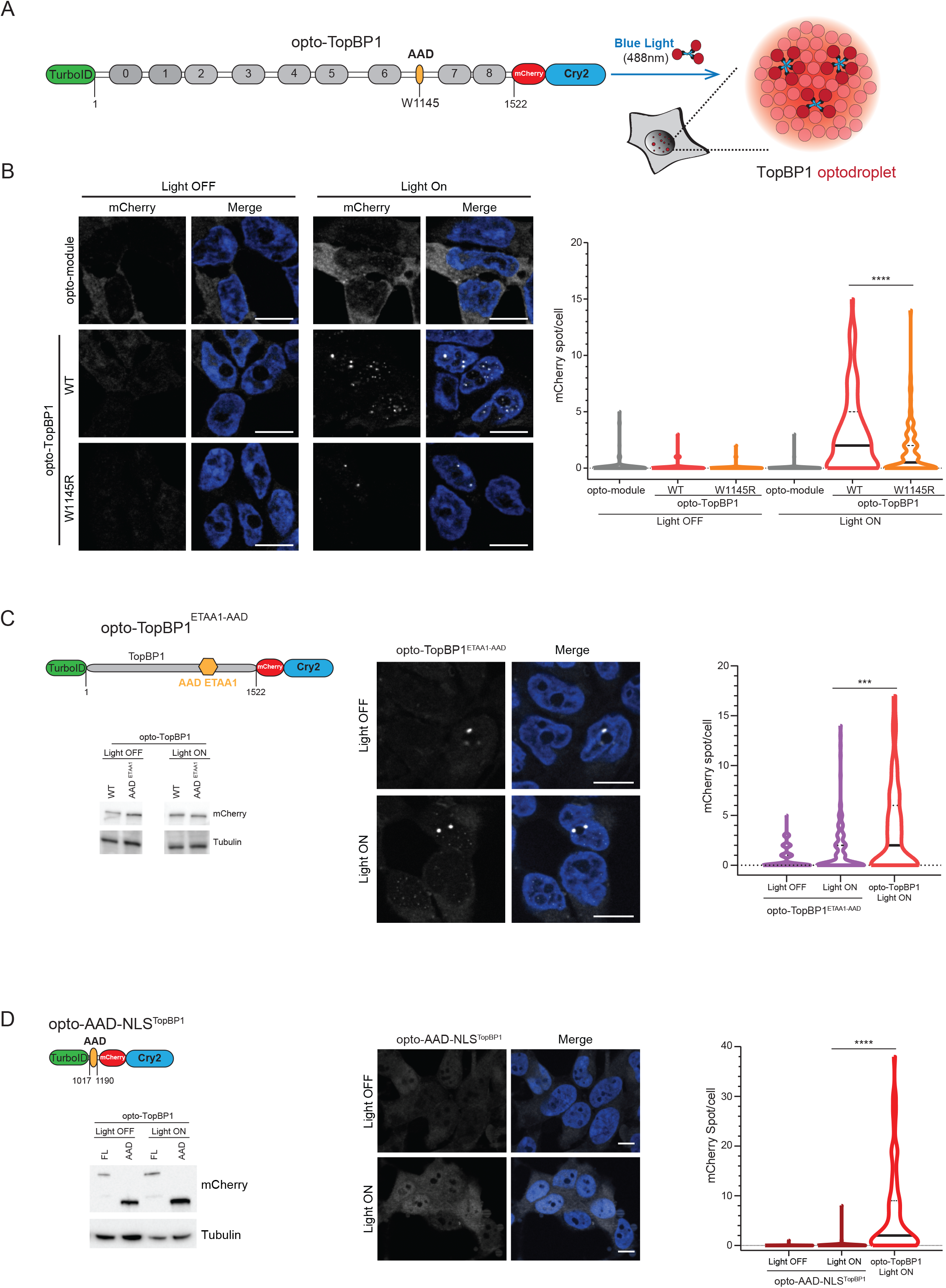
Optogenic control of TopBP1 biomolecular condensates. **A)** Schematic representation of the opto-TopBP1 construct. Full length WT or W1145R TopBP1 is fused at the N-terminus to TurboID, an optimized biotin ligase that can modify proteins within minutes, and at the C-terminus with mCherry fluorescent protein and the Cryptochrome 2 (Cry2) protein. Upon exposure to 488nm light, Cry2 oligomerises and induces TopBP1 condensation into mCherry positive foci. **B)** Representative fluorescence images of cells expressing WT and W1145R opto-TopBP1, before (Light OFF) and after (Light ON) 3min exposure to cycles of 4s light (488nm)-10s resting. Control cells express the opto-module mCherry-Cry2. **C-D)** Schematic representation of the opto-TopBP1 chimeric protein opto-TopBP1^ETAA1-AAD^ (**C**) and opto-AAD-NLS^TopBP1^ (**D**). TopBP1 ATR Activation Domain (AAD) was replaced by ETAA1 AAD (**C**) or cloned into the opto-module together with the TopBP1 Nuclear Localisation Sequence (NLS) (**D**). Representative fluorescence images of opto-TopBP1^ETAA1-AAD^ (**C**) and opto-AAD-NLS^TopBP1^ (**D**) exposed (Light ON) or not (Light OFF) to cycles of 4s light (488nm)-10s resting for 3min. Western blotting with the indicated antibodies showing opto-TopBP1, opto-TopBP1^ETAA1-AAD^ (**C**) and opto-AAD-NLS^TopBP1^ (**D**) expression. FL in **D** stands for Full Length and corresponds to opto-TopBP1 construct. **(B-C-D)** DNA was stained with Hoechst 33258. Scale bars: 10μm. Violin plot represents distribution of number of mCherry foci per cell. Median and quartile values are represented by continues and dashed lines respectively. The statistical significance of the difference in mCherry spot/cell distributions between samples is represented by *.

Besides TopBP1, Ewing’s tumor associated antigen 1 (ETAA1) has been recently identified as necessary to maintain Chk1 basal activity and stability (Michelena et al., 2019), and for the regulation of mitotic ATR signalling (Bass and Cortez, 2019). Like TopBP1, ETAA1 is endowed with an intrinsically disordered ATR activation domain (AAD) (Bass and Cortez, 2019; Bass et al., 2016; Haahr et al., 2016; Lee et al., 2016). Considering the conserved function and disordered nature of these ATR activation domains, we replaced the AAD of TopBP1 with the AAD of ETAA1, and then tested the capacity of the chimeric opto-TopBP1^ETAA1-AAD^ protein to form condensates. Exposure of these chimeric proteins to blue light did not yield condensates (Figure 1C). Consistent with this observation, ETAA1 did not form foci when fused to Cry2 and exposed to blue light (Suppl. Figure 1C). Furthermore, in isolation, the AAD of TopBP1 fused to Cry2 did not yield condensates upon optogenetic activation (Figure 1D). Collectively, these data indicate that TopBP1 can self-assemble into micron-sized condensates and that TopBP1 AAD has unique characteristic features that are necessary but not sufficient for TopBP1 higher-order assembly.

### Optogenetic and endogenous TopBP1 condensates share similar properties

To understand the nature of TopBP1 foci, we used the conceptual framework of biomolecular condensates to explore their properties. A three-dimensional analysis revealed that optogenetically-induced TopBP1 condensates have a spherical shape with an aspect ratio close to one, as if the interface of these structures was subjected to surface tension (Figure 2A + Suppl. video). Furthermore, opto-TopBP1 condensates occasionally fused, suggesting that TopBP1 undergoes dynamic clustering (Figure 2B). As expected for membraneless compartments that are held together by multiple weak and transient interactions, opto-TopBP1 foci were reversible (Figure 2C + Suppl. Figure 2A), yet dissolved within 15 minutes, while Cry2 oligomers disassemble spontaneously within 1-2 minutes (Shin et al., 2017). The relative stability of opto-TopBP1 condensates reflects the role of TopBP1 multivalent cooperative interactions in the formation of micron-sized protein assemblies. Next, we assessed the permeability of TopBP1 condensates to the surrounding milieu by fluorescence recovery after photobleaching, using an experimental system developed by Sokka et al., 2015. We induced expression of eGFP-TopBP1 in U-2-OS cells with doxycycline for 24 hours. In these experimental conditions, overexpressed eGFP-TopBP1 spontaneously forms foci that accumulate in nucleoli (Sokka et al., 2015). After photobleaching, eGFP-TopBP1 bodies recovered fluorescence signal within seconds, which reflects the rapid exchange of eGFP-TopBP1 molecules between the nucleoplasm and TopBP1 nuclear condensates (Suppl. Figure 2B). We conclude that the boundaries of TopBP1 compartments are permeable. We also noted that eGFP-TopBP1 condensates were sensitive to hexanediol, an aliphatic alcohol that disrupts weak hydrophobic interactions (Suppl. Figure 2C).

**Figure 2.**
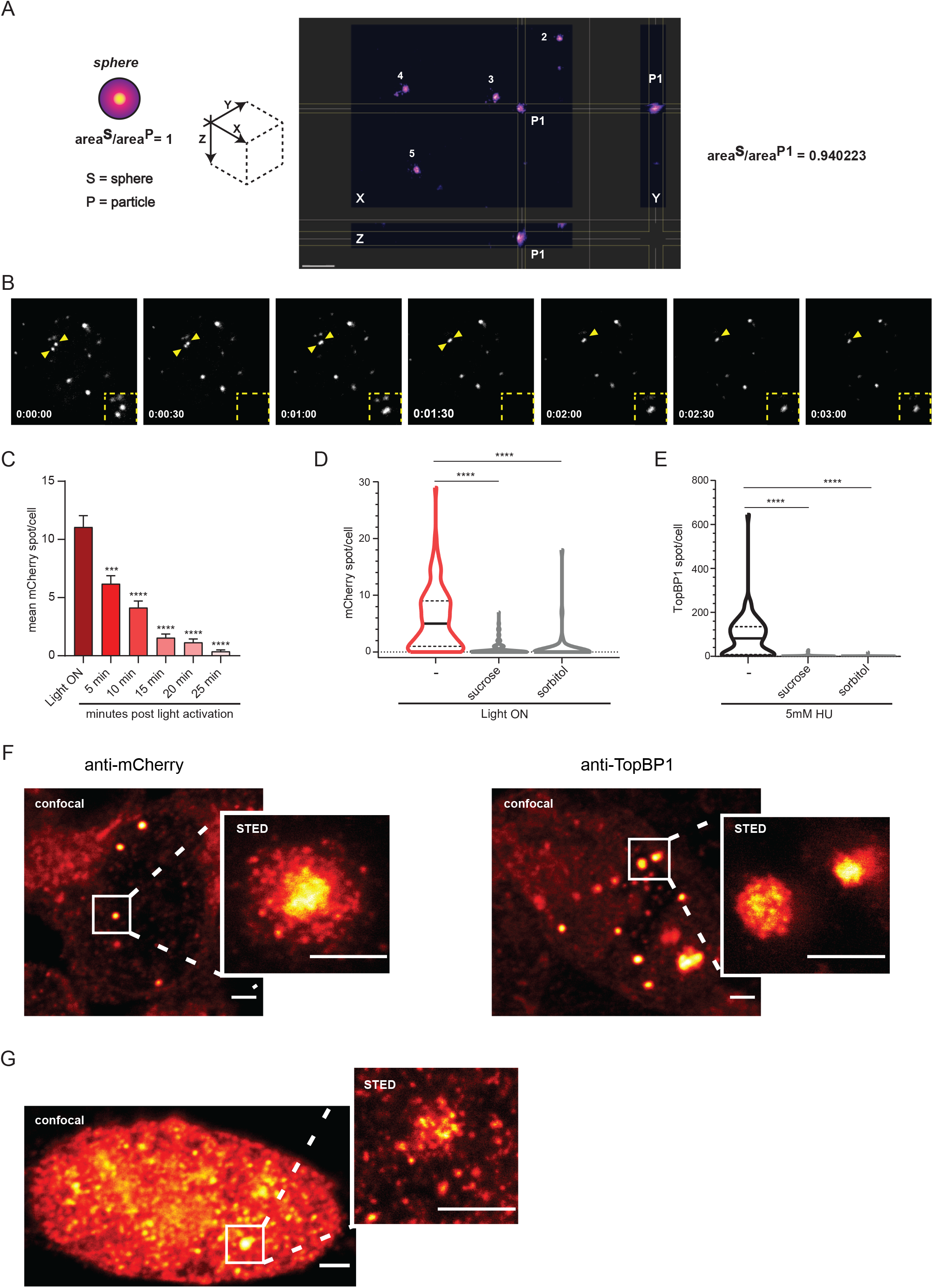
Properties of TopBP1 optogenetic and endogenous condensates. **A)** 3D analysis of opto-TopBP1 condensates by IMARIS software. To calculate sphericity: area^sphere^ /area^particle^. For a sphere the value is 1. We represented the 3D projection and the spherical value for particle 1 (P1). Other particles were analyzed for their spherical values (P2=0.932587; P3=0.918362; P4=0.949144; P5=0.935103). **B)** Fluorescence images representing a fusion event in cells expressing opto-TopBP1 WT activated by light (3min exposure to cycles of 4s light-10s resting). **C)** Histograms representing the mean of mCherry foci per cell. Cells expressing opto-TopBP1 WT were first exposed for 3min to cycles of 4s light (488nm)-10s resting (Light ON) and then left into dark from 5 to 25 min. **D)** Violin plot representing the number of mCherry foci per cell in cells expressing opto-TopBP1 WT after (Light ON) 3min exposure to cycles of 4s light (488nm)-10s resting. When indicated, cells were pre-treated with 0.5M sucrose or 0.4M sorbitol for 1h. **E)** Violin plot representing the number of endogenousTopBP1 foci per cell in cells treated with 5mM HU for 2h. When indicated, cells were simultaneously incubated with 0.5M sucrose or 0.4M sorbitol for 1h. Represented values correspond to the sum of three independent experiments. **D-E)** Median and quartile values are represented by continues and dashed lines respectively. The statistical significance of the difference in mCherry or TopBP1 foci/cell distributions between samples is represented by *. **F-G)** Super-resolution STED of opto-TopBP1 foci (**F**) and endogenous TopBP1 (**G**) foci identified with anti-mCherry (**F**-left panel) or anti-TopBP1 (**F**-right panel and **G**) antibodies. Opto-TopBP1 foci were induced by light (**F**), while endogenous TopBP1 foci by 5mM HU for 2h (**G**). Scale bars: 2μm for CONFOCAL images and 1μm for STED images.

The cooperative interactions that drive the formation of biomolecular condensates in physiological conditions are determined by protein sequences and by the properties of the surrounding milieu. Changes in osmotic concentration by addition of sorbitol or sucrose in the cell culture medium disrupt 53BP1 phase separation (Kilic, 2019). Likewise, osmotic stress inhibited both the assembly of TopBP1 condensates induced by optogenetic activation (Figure 2D + Suppl. Figure 2D), and the formation of endogenous TopBP1 foci induced by cellular treatment with the inhibitor of ribonucleotide reductase hydroxyurea (Figure 2E + Suppl. Figure 2E). This suggests that electrostatic forces drive the assembly of both synthetic (optogenetic) and endogenous TopBP1 condensates. Furthermore, recombinant opto-TopBP1 also formed nuclear foci when cells were exposed to hydroxyurea, rather than 488nm light (Suppl. Figure 2F), indicating that like endogenous TopBP1, opto-TopBP1 is mobilized and engages with endogenous components in response to stalled replication forks. TopBP1 condensates appear as homogenous structures by conventional fluorescence microscopy. We used stimulated emission depletion (STED) nanoscopy to gain better insights into the sub-structural organization of TopBP1 condensates. High-resolution imaging using an anti-mCherry antibody revealed the underlying sub-structure of optogenetic TopBP1 condensates, which consist in clusters of spherical, nanometer-sized particles (Figure 2F, left panel). TopBP1 clusters had a very tight appearance when detected with an anti-TopBP1 antibody that recognizes both endogenous and recombinant TopBP1 (Figure 2F, right panel). High-resolution imaging of endogenous TopBP1 foci induced by cellular treatment with hydroxyurea identified similar sub-structured clusters of nano-condensates (Figure 2G). Collectively, the data suggest that whether seeded by DNA replication impediments or Cry2 oligomerisation, the driving forces and the organization of TopBP1 nuclear condensates are similar.

### TopBP1 undergoes liquid-liquid phase separation *in vitro*

Multivalent protein scaffolds that drive the formation of biomolecular condensates often phase separate *in vitro*. To test if TopBP1 has the capacity to phase separate in physiologic salt and pH, we expressed the carboxy-terminal half of TopBP1 (amino acids 884-1522) in insect cells. This portion of TopBP1, hereafter referred to as b6-8, includes the BRCT 6 to 8 and the intrinsically disordered ATR activation domain (AAD), located between BRCT6 and BRCT7 (Figure 3A). TopBP1^b6-8^ mediates directly the activation of ATR, independently of its amino-terminal portion, which is required for the initiation of DNA replication (Hashimoto et al., 2006; Kumagai et al., 2006). TopBP1^b6-8^ formed optogenetically-induced foci in live cells (Suppl. Figure 3A). In the presence of the crowding agent Polyethylene Glycol (2%), purified TopBP1^b6-8^-GFP readily formed spherical condensates detected by fluorescence microscopy (Figure 3B, upper panel and Suppl. Figure 3B). Consistent with optogenetic experiments (Figure 1B), the W1145R substitution abolished the phase separation of TopBP1^b6-8^ (Figure 3B, lower panel). Addition of TopBP1^b6-8^-RFP to pre-formed TopBP1^b6-8^-GFP condensates yielded yellow compartments, suggesting that soluble TopBP1 molecules are recruited to TopBP1 condensates. By contrast, W1145R TopBP1^b6-8^-RFP did not stably associate with TopBP1^b6-8^-GFP condensates (Figure 3C). TopBP1 condensates were permeable to DNA, as revealed by the partitioning of double-strand DNA fragments into pre-formed TopBP1 condensates (Figure 3D). In the presence of a circular DNA plasmid (2.9 kb), TopBP1^b6-8^-GFP formed clusters of nano-condensates that were reminiscent, yet not equivalent, to high-resolution images of cellular TopBP1 condensates (Figure 3E). This observation suggests that the long anionic DNA polymer strongly influences the assembly of TopBP1 molecules. In isolation and in the absence of PEG, purified TopBP1^b6-8^-GFP (10μM) did not phase separate (Figure 3B minus PEG, and 3F). To recapitulate partially the complex environment of the nucleus, we spiked “Dignam and Roeder” nuclear extracts with recombinant TopBP1^b6-8^-GFP in physiologic salt concentration and pH. After incubation of the reaction mixture at 37° C, fluorescence microscopy analyses revealed μm-scaled TopBP1^b6-8^-GFP condensates (Figure 3F) co-localising with endogenous DNA (Suppl. Figure 3C). Altogether, the data suggest that TopBP1 foci-like structures are recapitulated with purified TopBP1 in the presence of a crowding agent or nuclear extracts. Consistent with optogenetic experiments in living cells, TopBP1^b6-8^-GFP condensation in nuclear extracts depends on the integrity of the AAD. We thus conclude that TopBP1, both *in vitro* and in living cells, has intrinsic capacity to form micron-sized condensates.

**Figure 3.**
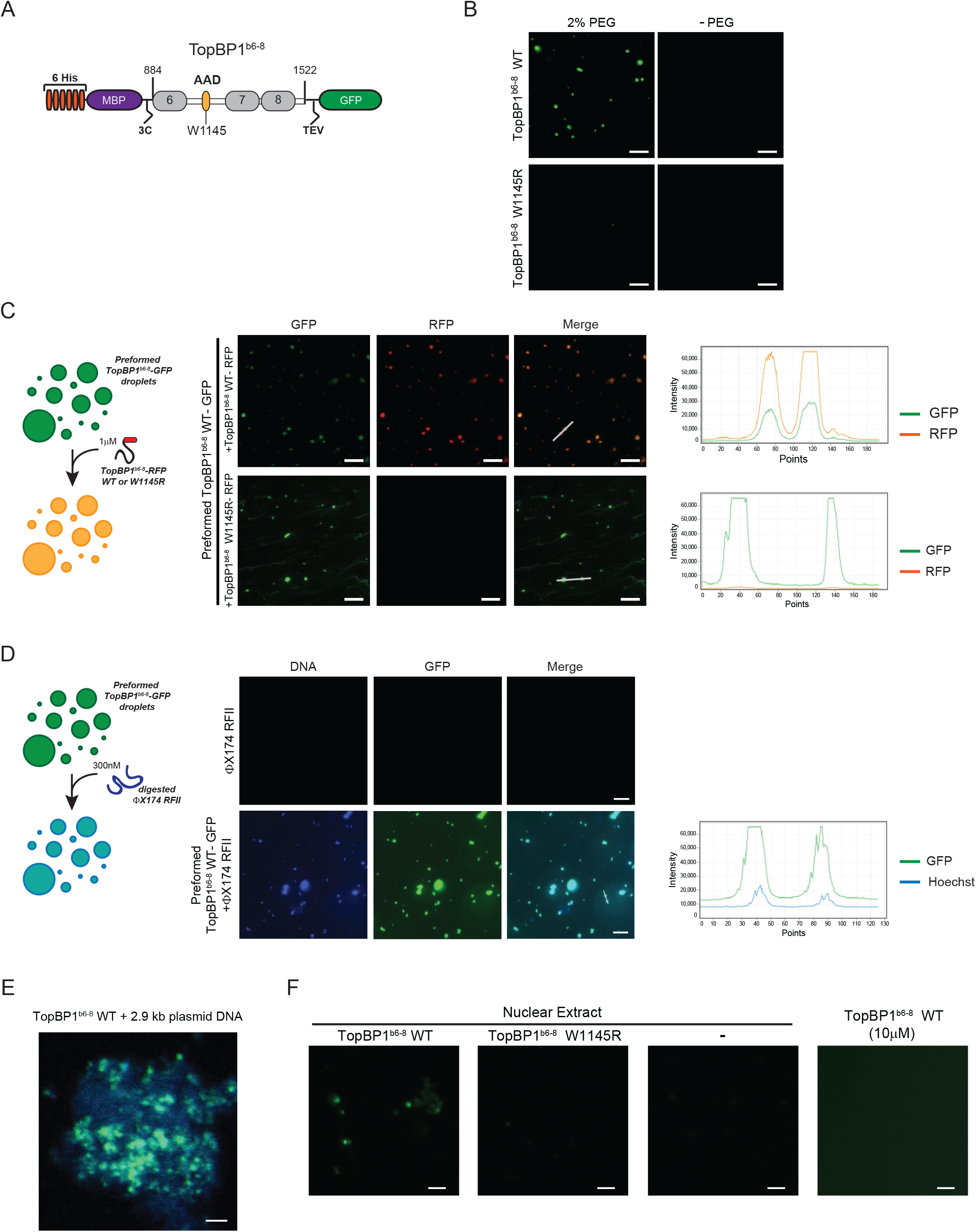
TopBP1^b6-8^ phase separates *in vitro*. **A)** Schematic representation of recombinant WT and W1145R TopBP1 (TopBP1^b6-8^) expressed in insect cells. The protein includes TopBP1 amino acids 884 to 1522, which comprise BRCT6-AAD-BRCT7-8. It is tagged with 6xHistidine and maltose binding protein (MBP) at its N-terminus, and monomeric Green Fluorescent Protein (GFP) at the C-terminus. PreScission (3C) and TEV protease sites are indicated. **B)** Representative images of TopBP1 droplets formed using 10µM of WT or W1145R TopBP1^b6-8^-GFP incubated in phosphate buffer pH 7.6 containing 150mM KAc and, when indicated, 2% PEG. Scale bars: 10μm. **C-D)** Representative images of TopBP1^b6-8^-RFP (1μM) (**C**) and double-stranded DNA (300nM) (**D**) trapping by pre-formed TopBP1^b6-8^-GFP (10μM) droplets in phosphate buffer pH 7.6, 150mM KAc, 2% PEG. The design of the experiment is represented schematically. DNA was stained with Hoechst 33258 (**D**). Scale bars: 10μm. Line scan of GFP-RFP (**C**) and GFP-Hoechst (**D**) signals is used to analyze co-localisation. **E)** Confocal image of purified (2.5µM) WT TopBP1^b6-8^ in the presence of 40ng 2.9 kb circular DNA plasmid DNA marked with Hoechst 33258. Scale bar: 2μm. **F)** Purified (2µM) WT and W1145R TopBP1^b6-8^ were incubated for 10min at 37°C with “Dignam and Roeder” nuclear extracts (0.2µg/µl), in physiological conditions (as described in Methods), and then analyzed by fluorescence microscopy for condensates in suspension. Control images of TopBP1^b6-8^-GFP (10μM) alone and of protein extract (0.2µg/µl) are shown. Scale bars: 10μm.

### Consequence of TopBP1 condensation on TopBP1-associated protein network

To test if recombinant TopBP1 condensates mimic endogenous TopBP1 foci, we analyzed the composition of TopBP1 condensates assembled in live cells. We used a biotin labelling approach with a low temporal resolution to gain a panoramic view of TopBP1 proximal proteins. A doxycycline-inducible cDNA encoding wild-type or W1145R TopBP1 fused to the mutated biotin ligase BirA* and the Flag epitope was stably integrated in Flp-In HEK293 cells (Figure 4A). We overexpressed Flag-BirA*-TopBP1 with doxycycline for 16 hours to a level that induces constitutive BirA*-TopBP1 condensates (Suppl. Figure 4), in the absence of DNA damaging agents. We labelled TopBP1 proximal proteins with biotin for 3 hours. More than 500 TopBP1 proximal proteins were identified by mass spectrometry with high confidence and reproducibility (Supplemental excel sheet). Proteins were ranked according to their iBAQ value (intensity-based absolute quantification), a proxy for protein abundance (Figure 4B). Among abundant TopBP1 proximal proteins, we identified known TopBP1 partners, including TOP2A, FANCJ, BRCA1, NBS1, MRE11, MDC1, 53BP1 and BLM (Figure 4B). Only five proteins showed differences in abundance between wild-type and W1145R TopBP1 (Log2 diff >2 with a p-value ≤ 0.05) (Figure 4C). Among them, only the nucleolar protein NOL11 was detected at high level, and, therefore, considered as a significant difference between wild-type and W1145R TopBP1. This observation is consistent with a previous report demonstrating that upregulated TopBP1 accumulates on ribosomal chromatin, segregates nucleolar components and yields nucleolar caps (Sokka et al., 2015). By contrast, W1145R substitution abrogates TopBP1 nucleolar segregation (Sokka et al., 2015). Consistent with this, W1145R TopBP1 had lost proximity with the nucleolar protein NOL11 (Figure 4D). Immunoblot analysis of TopBP1 proximal proteins confirmed that the substitution W1145R in the AAD of TopBP1 does not alter the proximity of TopBP1 with partner proteins implicated in ATR signalling, including ATR, BRCA1, MRE11 and FANCJ (Figure 4D). In sum, the substitution W1145R does not alter significantly the network of TopBP1 proximal proteins. In conclusion, at a low temporal resolution, whereas the composition of recombinant TopBP1 assemblies recapitulates endogenous TopBP1 foci, TopBP1 condensation does not increase significantly the association of TopBP1 with partner proteins.

**Figure 4.**
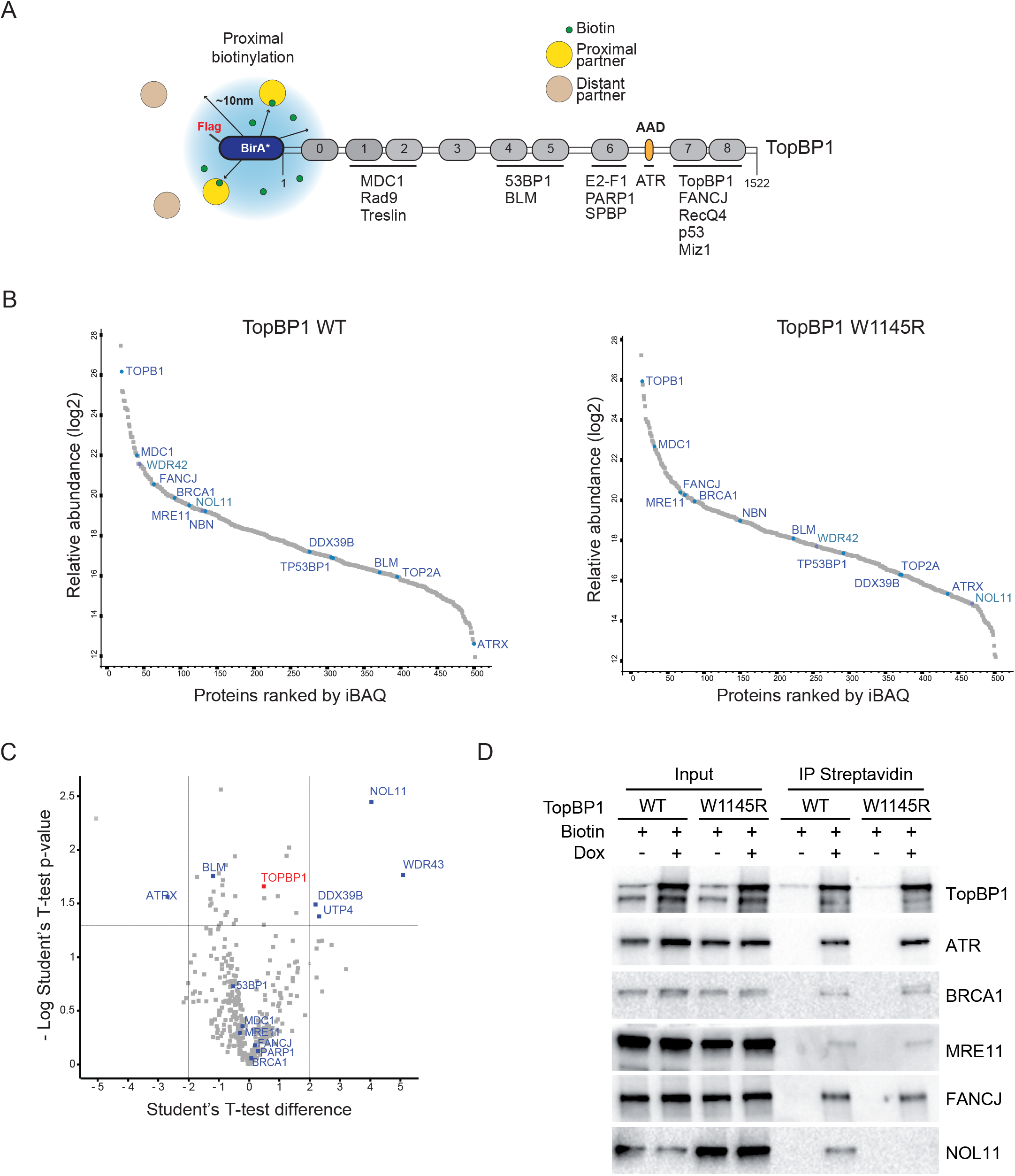
Reconstitution of TopBP1 protein network by BioID coupled to MS. **A)** Schematic representation of Flag-BirA*-TopBP1. In the presence of biotin (green dots), the biotin ligase BirA* biotinylates proteins within a 10nm range. **B-C)** Proteins identified by proximity-dependent biotin labelling in cells expressing WT or W1145R Flag-BirA*-TopBP1 were ranked according to their iBAQ value (**B**) or compared in a Volcano plot (**C**). Cells were grown for 16hr in presence of 1µg/ml doxycycline and incubated for 3h with 50µM biotin. For Volcano plot (**C**), a standard t-test was used to evaluate differences in protein abundance between samples. The left and right parts of volcano plot represent proteins enriched in W1145R and WT sample, respectively. Proteins depicted in the center part of the plot are common to both samples. **D)** Streptavidin pulldowns of proteins biotinylated by WT and W1145R Flag-BirA*-TopBP1 were probed for the indicated proteins by immunoblotting. When indicated (+), cells were treated with 1µg/ml doxycycline and 50µM biotin.

### Functional consequences of TopBP1 condensation

The optogenetic switch described above allows controlling TopBP1 assembly by light, which bypasses the confounding effects of prolonged cellular treatments with DNA damaging agents. In response to DNA replication impediments, ATR activates the effector checkpoint kinase 1 (Chk1) by phosphorylation on Ser345. To establish a functional link between TopBP1 condensation and ATR/Chk1 signalling, we exposed a cell culture dish to an array of blue-light LEDs for 3 minutes and then probed cell extracts for ATR mediated Chk1 phosphorylation on Ser345 by western blotting. Optogenetic condensation of TopBP1 induced robust phosphorylation of Chk1 Ser345 (Figure 5A) and of TopBP1 on Ser1138 (Figure 5B, t3), within 3 minutes, in the absence of an exogenous source of DNA damage. By contrast, the clustering defective W1145R TopBP1 mutant protein did not activate ATR/Chk1 signalling (Figure 5A). Consistent with the progressive dissolution of optogenetically-induced TopBP1 foci, phospho-Chk1 (Ser345) immunoblotting signals disappeared twenty minutes after optogenetic activation (Figure 5B, t23). Re-activation of TopBP1 condensates with blue light re-induced Chk1 phosphorylation (Figure 5B, t26), until dissolution of the reactivated TopBP1 condensates (Figure 5B, t46). We reiterated the process four times (Figure 5B). Collectively, the data indicate that Chk1 activation by ATR intersects precisely with TopBP1 condensation. This suggests that TopBP1 condensation operates as a switch-like mechanism that amplifies ATR activity above a threshold required for Chk1 phosphorylation.

**Figure 5.**
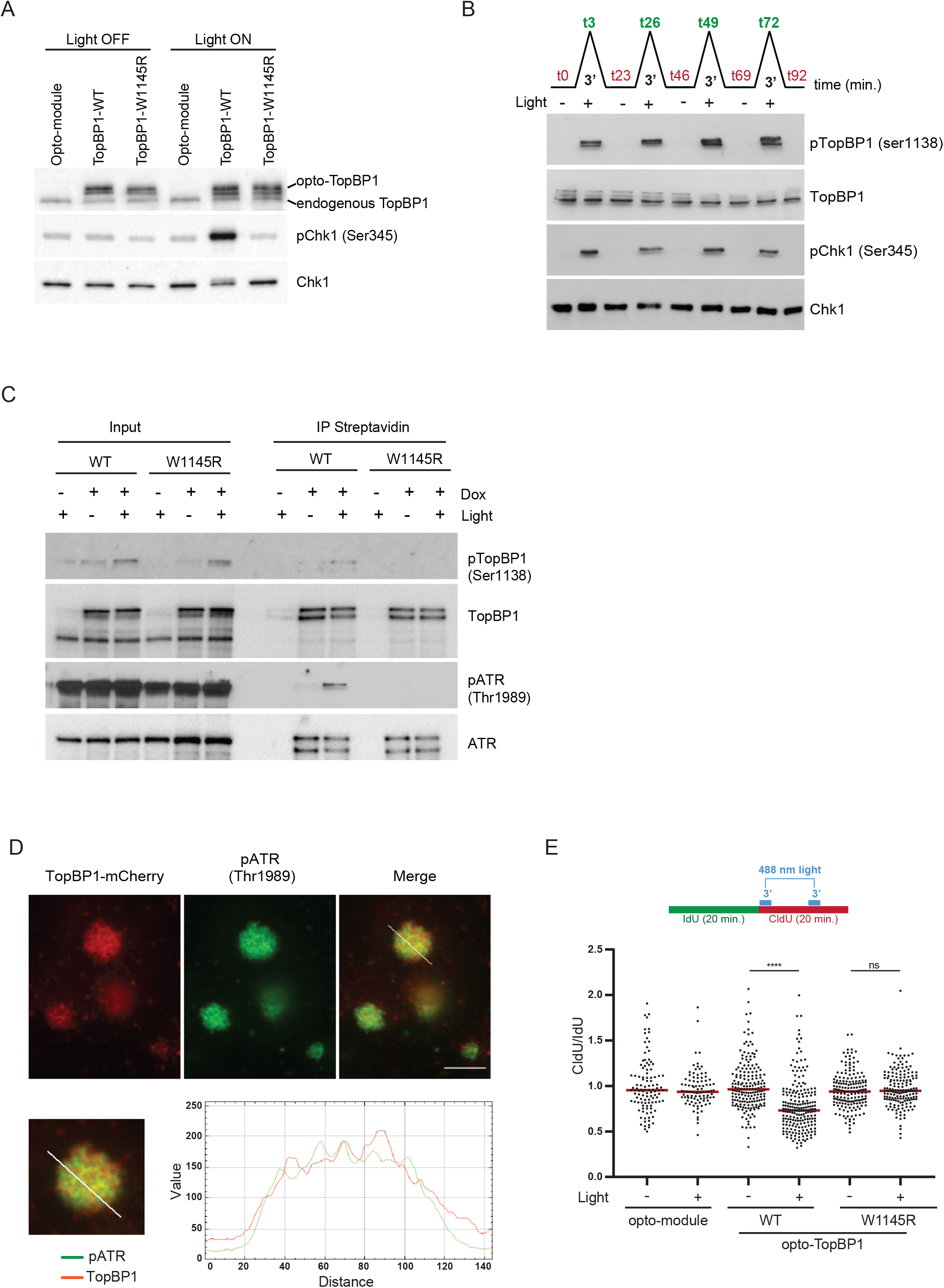
Functional consequences of TopBP1 condensation. **A)** Western blot analysis of TopBP1 and Chk1/phospho Chk1 (Ser345) of cells expressing WT and W1145R opto-TopBP1, before (Light OFF) and after (Light ON) 3min exposure to cycles of 4s light (488nm)-10s resting. Control cells express the opto-module mCherry-Cry2. **B)** Western blot analysis of TopBP1/pTopBP1 (Ser1138) and Chk1/pChk1 (Ser345) of cells expressing WT opto-TopBP1 during reiterative cycles of optogenetic activation (3min exposure to cycles of 4s light -10s resting). 4 cycles each 20 min. Total time: 92 min. **C)** Streptavidin pulldowns of proteins biotinylated by optogenetic activated WT and W1145R TurboID-TopBP1 were probed for the indicated proteins by immunoblotting. When indicated (+), cells were simultaneously incubated with 500µM of biotin and exposed to blue light for 10min of light-dark cycles (4s light followed by 30s dark). **D)** Super-resolution STED of opto-TopBP1 and pATR (Thr1989) foci. Co-localisation between opto-TopBP1 (anti-mCherry) and pATR is indicated by line scan. Scale bar: 1μm. **E)** CldU/IdU incorporation ratio of cells expressing WT and W1145R opto-TopBP1, before (-) and after (+) 3min exposure to cycles of 4s light (488nm)-10s resting. Mean values are represented by red lines and the statistical significance among them is represented by *. Control cells express the opto-module.

Of note, we observed the spontaneous activation of endogenous Chk1 in “Dignam and Roeder” nuclear extracts after 10 minutes incubation at 37° C (Suppl. Figure 5A). The level of phospho Chk1 (Ser345) signals was significantly higher than background signals observed when reaction mixtures were incubated at 4° C, or at 37° C in the presence of the ATR and mTOR inhibitor ETP46464 (Suppl. Figure 5A), confirming that the reaction was the product of ATR activity and occurred *in vitro*. In these experimental conditions, activation of endogenous Chk1 occurred in a purely endogenous protein extract. Neither recombinant TopBP1 protein nor exogenous ATR-activating DNA structures were added to the extract. Phosphorylation of Chk1 was blocked when nuclear extracts were pre-incubated with ethidium bromide, which disrupts protein-DNA interaction (Suppl. Figure 5B), indicating that endogenous DNA fragments present in the extracts are required for ATR activation. The spontaneous activation of endogenous ATR in nuclear extracts is compatible with the hypothesis that exceeding a critical protein concentration overcomes a kinetic barrier for the spontaneous assembly of functional condensates that activate ATR/Chk1 signalling.

In Figures 5A-B we showed that TopBP1 induced phosphorylation of Chk1 on Ser345. Once activated by ATR through Ser345 phosphorylation, Chk1 auto-phosphorylates on Ser296. This step is required for Chk1 to induce downstream molecular events leading to cell cycle arrest (Kasahara et al., 2010). Consistent with the hypothesis of opto-TopBP1 as functional assemblies, light-induced TopBP1 condensation yielded Chk1 Ser296 phospho-signals, confirming that Chk1 is active (Suppl. Figure 5C). UCN-01, an inhibitor of Chk1, blocked Chk1 auto-phosphorylation on Ser296, but had no major impact on Chk1 phosphorylation on Ser345 by ATR (Suppl. Figure 5 C). This confirms that the Chk1 phospho Ser296 signal is a product of Chk1 activity. To explore further the impact of TopBP1 condensation on TopBP1 interaction with partner proteins, we took advantage from the optogenetic tool shown in Figure 1A. The TopBP1-mCherry-Cry2 used for light-induced foci formation is tagged at its N-terminus with TurboID, an optimized biotin ligase that can biotinylated proteins within minutes (Branon et al., 2018). To detect ATR activity within optogenetic TopBP1 condensates, we induced TopBP1 condensation by blue-light illumination in the presence of biotin in the cell culture medium, and then purified biotinylated proteins with streptavidin-coated beads. We enriched phospho ATR (Thr1989) from cells expressing wild-type TopBP1 after optogenetic activation, specifically. We did not detect phospho ATR signals in proximity of the condensation-defective mutant W1145R TopBP1 (Figure 5C). In fluorescence microscopy, TopBP1 condensates induced by optogenetic activation co-localised with RAD9 and with phospho ATR (Suppl. Figure 5D-E). By contrast, Chk1 Ser345 phospho-signals rarely co-localised with TopBP1 condensates *per se*, but were detected in cells positive for TopBP1-mCherry condensates (Suppl. Figure 5F), consistent with Chk1 high mobility (Liu et al., 2006). Chk1 is not retained physically at DNA damage sites, allowing signal transmission from DNA damage sites to the rest of the cell (Liu et al., 2006). Super-resolution STED imaging revealed phospho ATR (Thr1989) signals intertwined with TopBP1 nano-condensates within TopBP1 clusters (Figure 5D). Collectively, these observations indicate that TopBP1 condensates function as reaction hubs.

As 100% of optogenetic TopBP1 condensates were chromatin bound (Suppl. Figure 5E), we analyzed the consequences of TopBP1 condensation on the progression of DNA replication forks using a DNA fiber labelling approach. We labelled DNA replication tracks with two consecutive pulses of the halogenated nucleotides IdU and CldU for 20 minutes each, and then visualized DNA replication tracks by fluorescence microscopy. We induced TopBP1 condensation during the CldU pulse using two cycles of 3 minutes blue-light illumination, in order to actuate and maintain TopBP1 foci during the 20 minutes labelling period (Figure 5E). DNA replication tracks labelled in the presence of optogenetic TopBP1 condensates were shorter than DNA replication tracks labelled in the absence of optogenetic activation. By contrast, blue-light illumination did not alter the progression of replication forks in cells that express the condensation defective mutant W1145R TopBP1. We conclude that TopBP1 condensates are functional entities. Mechanistically, TopBP1 condensation triggers activation of ATR/Chk1 signalling.

### Regulation of TopBP1 condensation

TopBP1 is among the proteins that are most highly phosphorylated and sumoylated in response to DNA replication stress (Munk et al., 2017). Post-translational modifications play a key role in the regulation of the molecular forces that drive the formation of biomolecular condensates (Soding et al., 2020). Since TopBP1 extensive capacity to self-assemble enables optogenetic induction of TopBP1 nuclear condensates, we used this optogenetic system to explore the role of phosphorylation in the formation of TopBP1 condensates. The activation of ATR/Chk1 signalling is dependent on TopBP1 phosphorylation (Burrows and Elledge, 2008; Hashimoto et al., 2006; Yoo et al., 2007). Thus, we reasoned that the basal kinase activity of ATR, which is independent of TopBP1 (Liu et al., 2011), may play a role in the condensation of TopBP1. To test this, we pre-incubated cells with the ATR inhibitors VE-821 (Charrier et al., 2011), or ETP-46464 (Llona-Minguez et al., 2014), for 1 hour, and then exposed opto-TopBP1 expressing cells to blue light for 3 minutes. In these experimental conditions, the formation of TopBP1 condensates was inhibited and the phosphorylation of Chk1 on the ATR site Ser345 was blocked (Figure 6A + Suppl. Figure 6A). To explore further the role of ATR in TopBP1 condensation, we mutated key amino acids in the AAD (Figure 6B). We substituted TopBP1 phenylalanine at position 1071 with alanine. This substitution locates within a predicted coiled coil in TopBP1 AAD and destabilized TopBP1 interaction with ATR (Thada and Cortez, 2019). The F1071A substitution abolished almost completely the optogenetic induction of TopBP1 condensates (Figure 6C), confirming that TopBP1 interaction with ATR is required for TopBP1 higher order assembly. In *Xenopus Laevis*, the phosphorylation of Ser1131 enhances the capacity of TopBP1 to activate ATR (Yoo et al., 2009). XTopBP1Ser1131 correspond to human Ser1138 TopBP1. The substitution Ser1138A inhibited light-induced TopBP1 condensation, whereas the phospho mimic Ser1138D stimulated TopBP1 condensation (Figure 6C). In blue-light exposed cells expressing F1071A TopBP1 or S1138A TopBP1, two mutations that impair TopBP1 condensation, the level of Chk1 phosphorylation on Ser345 was reduced in comparison with cells expressing wild-type TopBP1 (Figure 6C, right panel). Furthermore, phospho Ser1138 TopBP1 and phospho Thr1989 ATR signals were barely detectable in streptavidin pulldowns (Figure 6D). By contrast, the phosphomimetic substitution S1138D in TopBP1 yielded phospho Ser345 Chk1 signals upon optogenetic activation (Figure 6C, right panel) and phospho Thr1989 ATR was enriched in proximity of TopBP1 (Figure 6D). Collectively, the data suggest an amplification mechanism for activation of ATR/Chk1 signalling, whereby TopBP1 phosphorylation by ATR induces TopBP1 condensation, and TopBP1 condensation unleashes its capacity to activate ATR. We propose that TopBP1 condensation is a molecular switch that triggers checkpoint responses to DNA replication impediments.

**Figure 6.**
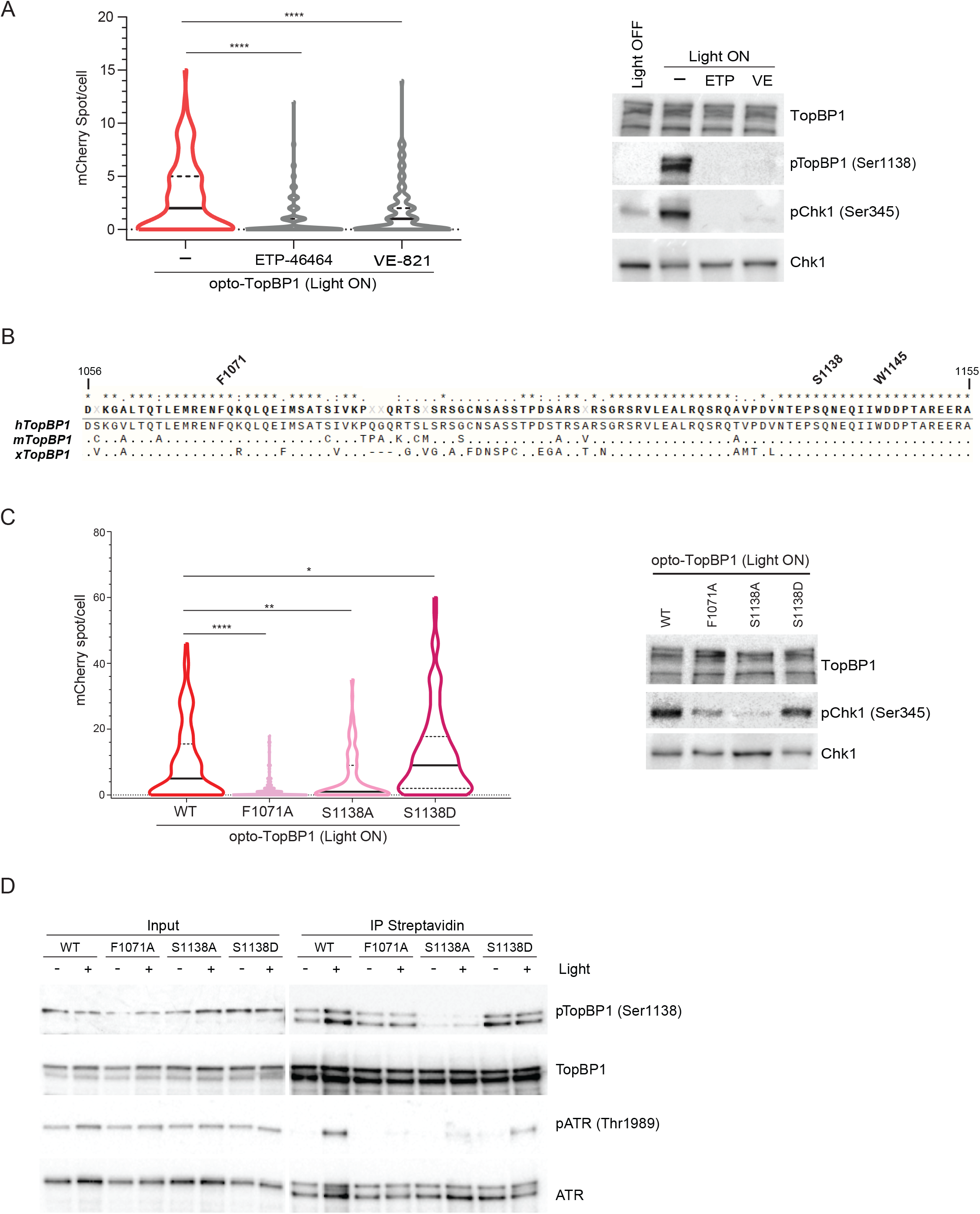
Regulation of TopBP1 condensation. **A)** (Left panel) Violin plot representing the number of mCherry foci per cell in cells expressing opto-TopBP1 WT after (Light ON) 3min exposure to cycles of 4s light (488nm)-10s resting. When indicated, cells were pre-treated with 10μM ATR inhibitors ETP-46464 or VE-821 for 1h. Median and quartile values are represented by continues and dashed lines respectively. (Right panel) Cells used for optogenetic experiments were probed for the indicated proteins by immunoblotting. The statistical significance of the difference in mCherry or TopBP1 foci/cell distributions between samples is represented by *. **B)** Alignment of ATR Activation Domain (AAD) sequences from Human (reference), Mouse and Xenopus. Amino acids from 1056 to 1155 of human TopBP1 are represented. Stars (*) on top of the alignment represent amino acids conserved among the three species. Conserved amino acids in mTopBP1 and xTopBP1 sequences are represented by single dots (.). Highly conserved amino acids F1071, S1138 and W1145 are indicated. **C)** (Left panel) Violin plot representing the number of mCherry foci per cell in cells expressing WT, F1071A, S1138A and S1138D opto-TopBP1 after (Light ON) 3min exposure to cycles of 4s light (488nm)-10s resting. Median and quartile values are represented by continues and dashed lines respectively. The statistical significance of the difference in mCherry or TopBP1 foci/cell distributions between WT and mutants samples is represented by *. (Right panel) Cells used for optogenetic experiments were probed for the indicated proteins by immunoblotting. **D)** Streptavidin pulldowns of proteins biotinylated by optogenetic activated WT, F1071A, S1138A and S1138D TurboID-TopBP1 were probed for the indicated proteins by immunoblotting. When indicated (+), cells were simultaneously incubated with 500µM of biotin and exposed to blue light for 10min of light-dark cycles (4s light followed by 30s dark).

## Discussion

In this study, we provide evidence that TopBP1 is a protein scaffold that can self-assemble extensively to yield tight clusters of nano-condensates. TopBP1 condensation functions as a molecular switch for ATR/Chk1 signalling. We show that TopBP1 condensation depends on TopBP1 interaction with ATR, on the basal kinase activity of ATR and on the phosphorylation of TopBP1. We obtained evidence that the phosphorylation of TopBP1 on serine 1138 is essential for TopBP1 condensation. Thus, we propose a refined model of ATR activation (Figure 7): TopBP1 association with ATR yields a positive feedback loop, with TopBP1 condensation acting as a mechanism of amplification promoting ATR signal transduction. In the early stages of ATR signalling, ATR-ATRIP and TopBP1 congregates on RPA-coated single-stranded DNA, the 9-1-1 complex is loaded at single to double strand DNA junctions and stabilizes TopBP1. In later stages, the phosphorylation of TopBP1 induces its higher-order assembly into micron-sized clusters of nano-condensates, where ATR activity is amplified and the checkpoint signal transmitted. Our combined approach enabling the concomitant induction of TopBP1 condensation and the labelling of TopBP1 proximal proteins confirms that TopBP1 condensates are the sites of ATR activation. The yeast homologue of ATR, Mec1, has been proposed to be activated via an allosteric mechanism (Wang et al., 2017). We surmise that the compartmentation of ATR signalling proteins creates a reaction hub where the probability of molecular interactions required for ATR activation is increased. A TopBP1 clustering mechanism for ATR activation is reminiscent of Ras nano-clusters, which are the sites of Ras effector recruitment and activation. Ras clusters assemble transiently on the plasma membrane, function as high-gain amplifiers and are critical for MAPK signal transduction (Prior et al., 2003; Tian et al., 2007).

**Figure 7.**
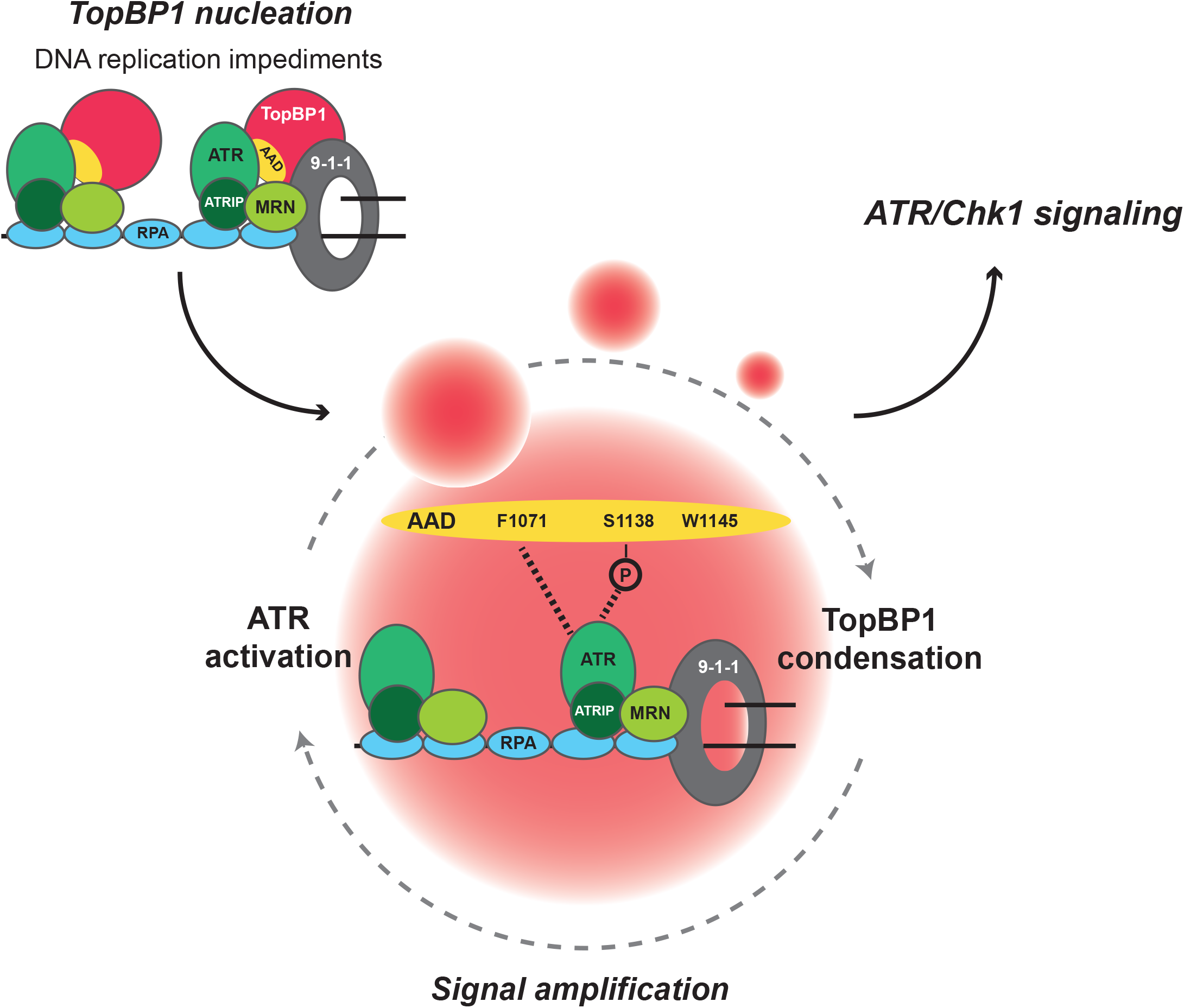
TopBP1 condensation switches on ATR/Chk1 signalling. Model showing the accumulation of ATR/ATRIP and TopBP1 on RPA coated single-stranded DNA. Interaction with ATR through the ATR Activation Domain (AAD), phosphorylation of TopBP1 and ATR kinase activity induce TopBP1 condensation. The enrichment of molecular reactants in TopBP1 compartments stimulates ATR activity above a threshold required for Chk1 activation and signal amplification.

The data shown here suggest that the molecular forces driving TopBP1 condensation are multiple weak and highly cooperative interactions of TopBP1 molecules. First, whether seeded by hydroxyurea-induced replication stress or Cry2 oligomerisation, TopBP1 nuclear condensates are dissolved upon addition of sorbitol or sucrose in the cell culture medium. These compounds destabilize weak electrostatic interactions involved in protein phase separation. The aliphatic alcohol hexanediol also dissolves TopBP1 condensates, suggesting that hydrophobic interactions contribute to TopBP1 higher order assembly. Second, purified TopBP1 undergoes liquid-liquid phase separation in physiologic salt and pH, a characteristic feature of multivalent protein scaffolds that underpin the formation of membrane-less compartments. It is noteworthy that TopBP1 phase separation *in vitro* occurred not only in the presence of the crowding agent PEG, but also in the complex environment of a nuclear extract, where multiple homotypic and heterotypic interactions could influence the capacity of TopBP1 to self-assemble. Third, TopBP1 condensation was highly sensitive to key amino acid substitutions and post-translational modifications within its intrinsically disordered ATR activation domain. These modifications typically change the cooperative molecular forces that organize protein condensation. Point mutations in the AAD that locally change the charge (W1145R, S1138D), the hydrophobicity (F1071A) or the polarity (S1138A) modulate the growth of TopBP1 condensates induced by optogenetic activation. Furthermore, the substitution W1145R in the AAD impaired the phase separation of purified TopBP1^b-6-8^ *in vitro*. Last, our data indicate that the integrity of the TopBP1 AAD is essential but not sufficient for the growth of TopBP1 nuclear condensates. In isolation, the AAD of TopBP1 fused to Cry2 did not yield condensates, while the carboxy-terminal half of TopBP1, which includes the AAD and BRCT6-8, assembled condensates. This indicates that the BRCT6-8 also contributes to TopBP1 higher-order assembly, consistent with data showing that BRCT7/8 promotes TopBP1 oligomerisation (Chowdhury et al., 2014).

The assembly of complex heterogeneous biomolecules via multivalent, weak, dynamic and cooperative interactions is emerging as a common principle of compartmentation in the nucleus (Banani et al., 2017; Shin and Brangwynne, 2017; Soding et al., 2020). We expect the rich diversity of molecular associations in live cells to yield diverse types of condensates with distinct composition and biophysical properties. TopBP1 is a scaffold protein involved in different pathways essentials for cell survival. In this study, we focused on its role in ATR activation. TopBP1 condensates induced by optogenetic activation were variable in size and fused occasionally, two characteristic features of non-stoichiometrically defined biomolecular condensates. FRAP analysis indicates that the exchange of TopBP1 molecules outside and inside TopBP1 compartments was unimpeded. High-resolution imaging by stimulated emission depletion (STED) microscopy revealed that TopBP1 micron-sized foci detected by conventional microscopy are made of clusters of nano-condensates, an organization that is likely to reflect the role of chromatin in TopBP1 higher-order assembly. Consistent with this, our *in vitro* studies suggest that long DNA polymers have major influence on the condensation of purified TopBP1. The organization of TopBP1 into tight clusters of nano-condensates was common to both recombinant TopBP1 condensates induced by optogenetic activation and endogenous TopBP1 condensates induced by DNA replication impediments. Interestingly, in mitosis, TopBP1 forms filamentous structures that bridge MDC1 foci (Leimbacher et al., 2019), suggesting that specific molecular associations dictate the organization of TopBP1 higher-order structures. The sub-structural organization of TopBP1 condensates is reminiscent of 53BP1 foci, which consist in nano-domains assembled into macro-domains (Ochs et al., 2019). Single-molecule localisation microscopy indicate that a 53BP1 repair focus is a composition of multiple clusters of 53BP1 molecules (McSwiggen et al., 2019), and its internal architecture is determined both by DNA topology and protein-protein interactions that provide a structural scaffold (Ochs et al., 2019).

The function of TopBP1 condensation described here explains previous observations. In *Saccharomyces cerevisiae*, artificial co-localisation of the 9-1-1 complex and Ddc2^ATRIP^-Mec1^ATR^ via tethering to an array of 256 LacO repeats bypasses the requirement for DNA damage to activate Mec1^ATR^ (Bonilla et al., 2008). In *Schizosaccharomyces pombe*, artificial tethering of either one of Rad3^ATR^, RAD4^TopBP1^ or RAD9 to a LacO array triggers a checkpoint response that utilizes the endogenous proteins (Lin et al., 2012). The LacO recruited Rad3^ATR^ must phosphorylate endogenous RAD9 to promote Rad4^TopBP1^ recruitment and activate the checkpoint (Lin et al., 2012). Furthermore, TopBP1 activates ATR *in vitro* and in cells when artificially tethered to DNA (Lindsey-Boltz and Sancar, 2011). Based on the findings described here, we surmise that the artificial tethering of checkpoint proteins to LacO arrays is nucleating the condensation of endogenous proteins, which switches on checkpoint signalling.

The transient and reversible nature of the molecular forces that underpin the formation of functional TopBP1 nuclear condensates appears well adapted to cellular regulation and optimal responsivity to DNA replication impediments, as opposed to the stable interaction of proteins that characterize molecular machines with defined stoichiometry. The formation of functional micron-sized condensates through the regulated self-assembly of multivalent protein scaffolds may represent a fundamental principle underlying the formation of functional nuclear foci in response to DNA damage.

## Supporting information

TopBP1 BioID proteomics

sphericity TopBP1

## Acknowledgments

We thank all members of the laboratory, Olivier Ganier and Pierre-Henri Gaillard for their critical reading of the manuscript. We are grateful to Lee Zou for the cDNA encoding TopBP1, Brian Raught for pCDNA5_FRT-TO_FlagBirA*, Clifford P Brangwynne for the cDNA encoding Cry2, Juhani E. Syväoja for the U-2-OS cell lines expressing eGFP-TopBP1 and Simon Alberti for technical information. This work was supported by MSD Avenir, by la Fondation ARC pour la recherche sur le cancer (PGA1 RF20180206787), and by the SIRIC Montpellier Cancer (grant INCa_Inserm_DGOS_12553).

## Author contribution

Conceptualization: C.F., J.B. and A.C; Methodology: C.F., J.B. and A.C.; Investigation: C.F., A.P., S.V., E.A., M.L., M-P.B., S.U. and J.B.; Data curation: C.F., A.P. J.B and A.C. Writing-original draft: C.F., J.B. and A.C. Writing-Review & Editing: C.F., A.P., J.B. and A.C. Supervision: J.B. and A.C. Project Administration: A.C. Funding Acquisition: A.C.

## Methods

### Resources tables

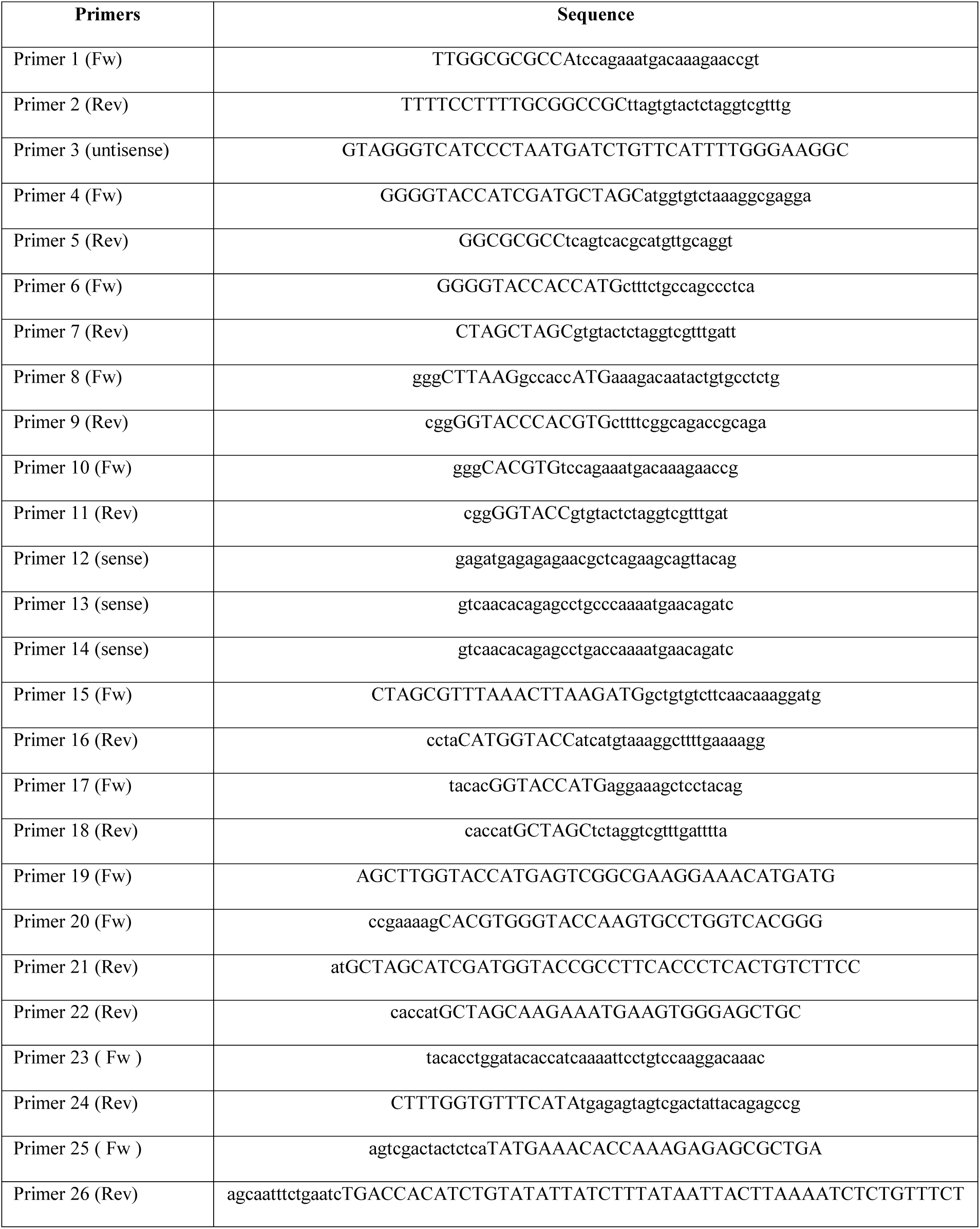
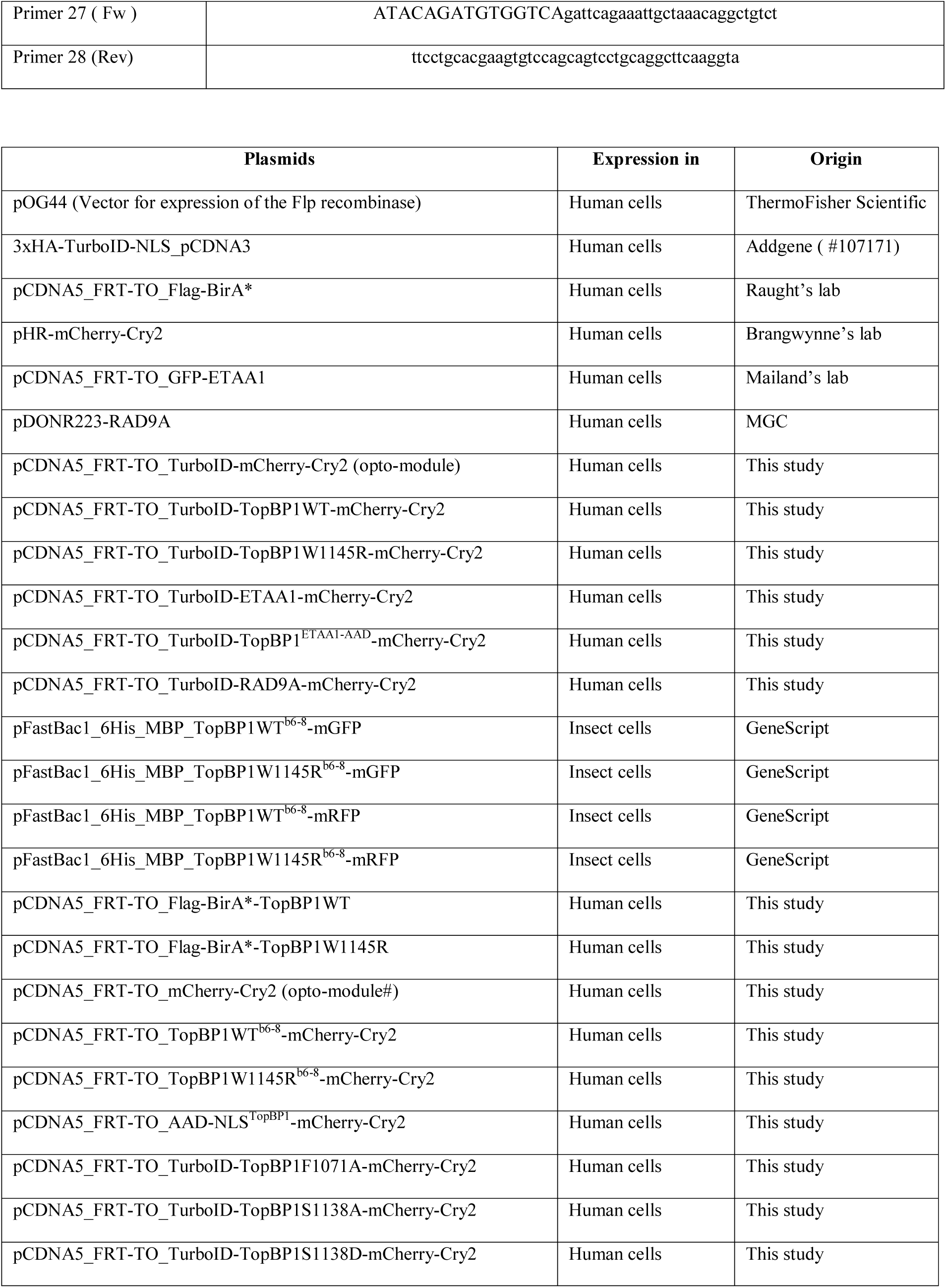

### Antibodies

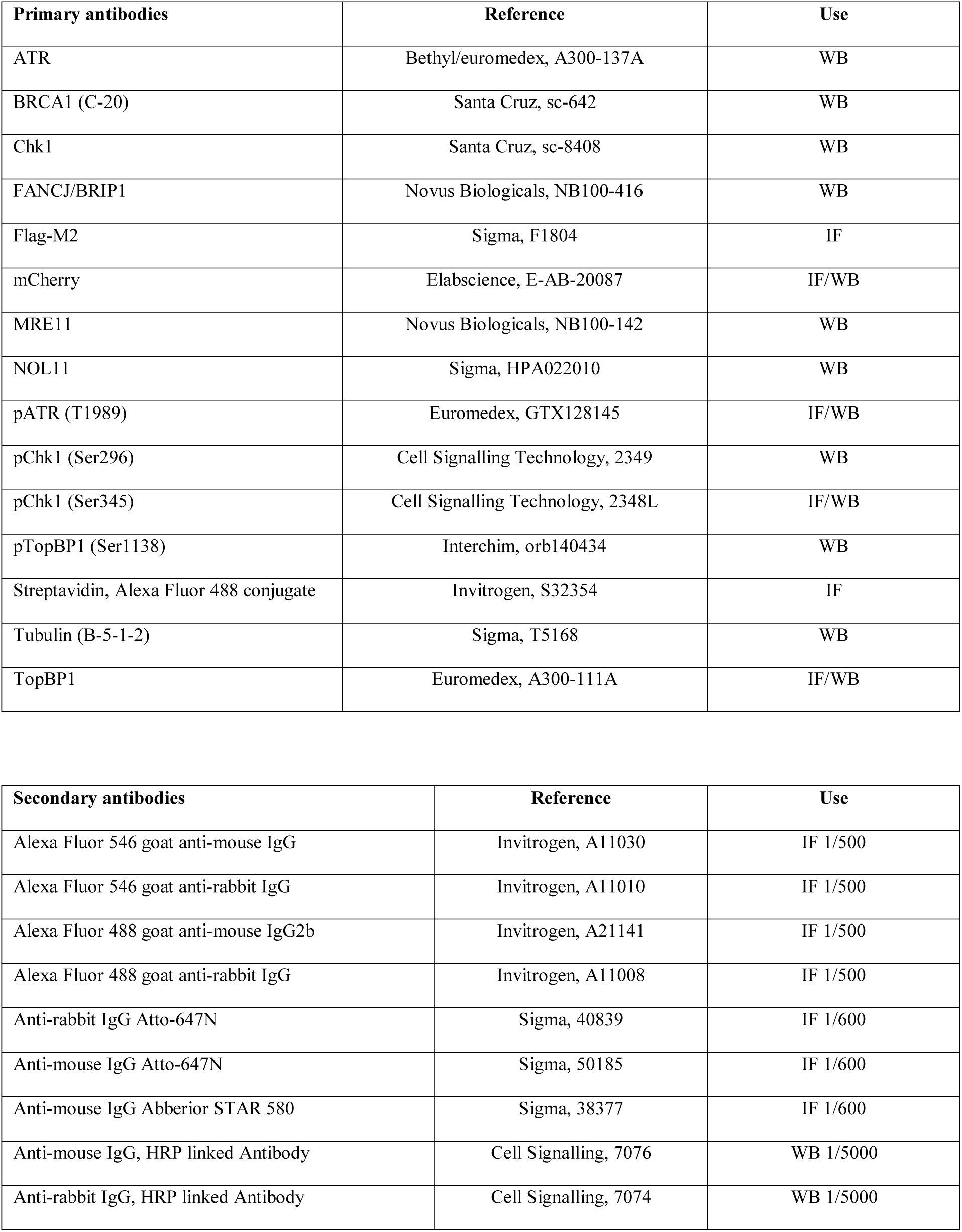

### Cell lines

*<u>Flp-In™T-REx™293</u>* and *<u>HEK293</u>* cell lines were grown under standard conditions (37°C, 5% CO_2_) in Dulbecco’s modified Eagle’s medium (Merck-Sigma-Aldrich, D5796). For *Flp-In™T-REx™293* the medium was supplemented with 10% fetal bovine serum (FBS), 100µg/ml Zeocin and 15µg/ml Blasticidin. *Flp-In™T-REx™293* transfected cells were selected and maintained with 15µg/mL Blasticidin and 150µg/mL Hygromycin.

*<u>U-2-OS Tet-On</u>* cell lines expressing eGFP-TopBP1 WT were grown under standard conditions (37°C, 5% CO_2_) in modified McCoy’s 5a medium (Merck-Sigma-Aldrich, M9309) supplemented with 10% fetal bovine serum (FBS), 100μg/ml of Hygromycin and 200μg/ml of G418 as selective antibiotics (Sokka et al., 2015). Expression of eGFP-TopBP1 WT was induced with 1μg/ml of doxycycline for 24 hours.

### Sf9 insect cells culture conditions

Sf9 cells were grown in EX-CELL® 420 Serum-free medium (Sigma-Aldrich, 14420C). Cells were maintained between 2×10^6^ and 1×10^7^ cells/ml at 28°C in flasks (agitation 140 rpm).

### Plasmid constructs

For pCDNA5_FRT-TO_Flag-BirA*-TopBP1 WT or W1145R, TopBP1 full-length cDNA (a kind gift from Lee Zou) was amplified by PCR with primers 1 and 2 using Phusion® High-Fidelity DNA Polymerase (New England Biolabs, CM0530). The forward and reverse primers contain *AscI* and *NotI* sites, respectively. The amplified PCR was inserted into the pCDNA5_FRT-TO_Flag-BirA* (a kind gift from Biran Raught) linearised with *AscI*/*NotI* digestion.

For pCDNA5_FRT-TO_mCherry-Cry2 (opto-module#), the Flag-BirA* fragment was deleted from the pCDNA5_FRT-TO_FlagBirA* using *KpnI/AscI* enzymes and replaced with the mCherry-Cry2 fragment amplified by PCR from the plasmid pHR-mCherry-Cry2 (a kind gift from Brangwynne’s lab) with primers 4 and 5. In the second step, TopBP1 amino acids 884 to 1522 fragment (BRCT6-AAD-BRCT7-8 WT or W1145R) was amplified with primers 6 and 7, digested with *KpnI/NheI* and inserted into pCDNA5_FRT-TO_mCherry-Cry2 to produce opto*-*TopBP1^b6-8^ WT and opto*-*TopBP1^b6-8^ W1145R. For pCDNA5_FRT-TO_TurboID-mCherry-Cry2 construction (opto-module), the TurboID fragment was amplified by PCR from the 3xHA-TurboID-NLS_pCDNA3 plasmid (Addgene #107171) with primers 8 and 9, digested with *AflII*/*KpnI* enzymes and inserted into the opto-module# to produce the opto-module. In the second step, TopBP1 WT and W1145R were amplified by PCR with primers 10 and 11, digested with *PmlI/KpnI* enzymes and inserted into the opto-module construct. Mutations in the AAD of TopBP1 were generated using the “*QuickChange Multi Site-directed mutagenesis kit*” (Agilent technologies, C200515): W1145R with primer 3, F1071A with primer 12, S1138A with primer 13, S1138D with primer 14. pCDNA5_FRT-TO_TurboID-AAD-NLS^TopBP1^-mCherry-Cry2 was generated by PCR amplification of TopBP1 AAD (primers 15-16) and TopBP1 NLS (primers 17-18). Fragments were inserted in 2 steps into the opto-module digested with *AflII*/*KpnI* (TopBP1 AAD) and *KpnI/NheI* (TopBP1 NLS) enzymes respectively. pCDNA5_FRT-TO_TurboID-ETAA1-mCherry-Cry2 was generated by PCR amplification of ETAA1 with primers 19 and 20 on pCDNA5_FRT-TO _GFP-ETAA1 (a kind gift from Mailand’s lab) and insertion of the fragments into the opto-module digested with *KpnI/NheI* enzymes. pCDNA5_FRT-TO_TurboID-RAD9A-mCherry-Cry2 was generated by PCR amplification of RAD9 with primers 21 and 22 on pDONR223-RAD9A (obtained through MGC Montpellier Genetic Collections) and fragment was cloned into the *KpnI*-digested opto-module following the *In-Fusion HD Cloning Kit* protocol. The plasmid with the chimeric constructs of TopBP1 carrying the AAD of ETAA1 (pCDNA5_FRT-TO_TurboID-TopBP1^ETAA1-AAD^-mCherry-Cry2) was generated using the *NEBuilder HiFi DNA assembly Master Mix* (New England Biolabs, E2621L). This kit was used to assemble multiple DNA fragments with 30 bp-overlap and replace 1530 bp inside TopBP1. DNA fragments were produced by PCR: oligos 23-24 were used to amplify TopBP1 sequence before the AAD (PCR on any plasmid containing full-length TopBP1), oligos 25-26 were specific for the AAD of ETAA1 (PCR on pCDNA5_FRT-TO _GFP-ETAA1) and oligos 27-28 were specific for TopBP1 portion after the AAD (PCR on any plasmid containing full-length TopBP1). To obtain the chimeric constructs, PCR products were assembled according to the manufacturer’s instruction and ligated into pCDNA5_FRT-TO_TurboID-TopBP1-mCherry-Cry2 digested with *EcoNI/SbfI*.

pFastBac1 plasmids containing WT and W1145R BRCT 6 to 8 fragments of TopBP1 were synthesized by GeneScript after codon optimization for expression in insect cells (sequence available upon request), and sub-cloned into the 6His-MBP_3C_MCS_TEV_mRFP and 6His-MBP_3C_MCS_TEV_mGFP cassette using the restriction sites *EcoRI/KpnI*.

### Western Blotting

Whole cell extracts were lysed with 1X Laemmli Sample buffer (Biorad, C161-0737) and heated 5min at 95°C. Cell extracts were resolved using pre-cast SDS-PAGE (7.5% and 10%) from BioRad and transferred to nitrocellulose membrane using a transfer apparatus according to the manufacturer’s instructions (BioRad). Membranes were saturated with 10% non-fat milk diluted in TBS-0.2% Tween 20 (TBS-T), incubated with primary antibodies overnight at 4°C and with anti-mouse-HRP or anti-rabbit-HRP secondary antibodies for 1h. Blots were developed with ECL according to the manufacturer’s instructions.

### Affinity capture of biotinylated proteins: BioID

*Flp-In™ T-REx™ 293* cell lines stably transfected with Flag-BirA*-TopBP1 WT or W1145R grown to 75% confluence were incubated with 1µg/ml of doxycycline (Clontech, 631311) for 16h and with 50µM biotin for 3 or 16 hours. Cells were washed with PBS and lysed with lysis buffer (50mM Tris-HCl pH 7.5, 150mM NaCl, 1mM EDTA, 1mM EGTA, 1% NP-40, 0.2% SDS, 0.5% Sodium deoxycholate) supplemented with 1X complete protease inhibitor (Roche) and 250U benzonase (Sigma, CE1014). Lysed cells were incubated on a rotating wheel for 1h at 4°C prior sonication on ice (40% amplitude, 3 cycles 10sec sonication-2sec resting). After 30min centrifugation (7750 rcf) at 4°C, the cleared supernatant was transferred to a new tube and total protein concentration was determined by Bradford protein assay (BioRad, C500-0205). For each condition, 300µg of proteins were incubated with 30µl of Streptavidin-Agarose beads (Sigma, CS1638) on a rotating wheel at 4°C for 3hr. After 1min centrifugation (400 rcf), beads were washed, successively, with 1ml of lysis buffer, 1ml wash buffer 1 (2% SDS in H_2_O), 1ml wash buffer 2 (0.2% sodium deoxycholate, 1% Triton X-100, 500mM NaCl, 1mM EDTA, and 50mM Hepes pH 7.5), 1ml wash buffer 3 (250mM LiCl, 0.5% NP-40, 0.5% sodium deoxycholate, 1mM EDTA, 500mM NaCl and 10mM Tris pH 8) and 1ml wash buffer 4 (50mM Tris pH 7.5 and 50mM NaCl). Bound proteins were eluted from the magnetic beads using 80µl of 2X Laemmli Sample buffer and incubated at 95°C for 10min. 10% of the sample was used for Western blot analysis. For the *Flp-In™ T-REx™ 293* cell lines stably transfected with the doxycycline-inducible TurboID-TopBP1WT-mCherry-Cry2 or the mutated forms of TopBP1, cells were simultaneously incubated with 500µM of biotin and exposed to blue light for 10min of light-dark cycles (4s light followed by 30s dark). Biotin proximity labelling of light-induced TopBP1 partners were pulled-down using streptavidin-coated beads as described before and probed by immunoblotting to detect proteins that are associated with TopBP1 clusters, in absence of DNA damage.

### Mass spectrometry

Sample digestion was essentially performed as described (Shevchenko et al., 2006). Briefly, proteins were loaded on a SDS-PAGE (BioRad, 456-1034) and, after short migration, a single band was excised. Proteins in the excised band were digested with Trypsin (Promega, V5280). The resulting peptides were analyzed online by nano-flow HPLC-nanoelectrospray ionization using a Qexactive HFX mass spectrometer (Thermo Fisher Scientific) coupled to a nano-LC system (Thermo Fisher Scientific, U3000-RSLC). Desalting and preconcentration of samples were performed online on a Pepmap® precolumn (0.3 x 10mm; Fisher Scientific, 164568). A gradient consisting of 0% to 40% B in A (A: 0.1% formic acid [Fisher Scientific, A117], 6% acetonitrile [Fisher Scientific, A955], in H_2_O [Fisher Scientific, W6], and B: 0.1% formic acid in 80% acetonitrile) for 120min at 300nl/min was used to elute peptides from the capillary reverse-phase column (0.075 x 250mm, Pepmap®, Fisher Scientific, 164941). Data were acquired using the Xcalibur software (version 4.0). A cycle of one full-scan mass spectrum (375–1,500m/z) at a resolution of 60000 (at 200m/z) followed by 12 data-dependent MS/MS spectra (at a resolution of 30000, isolation window 1.2m/z) was repeated continuously throughout the nanoLC separation. Raw data analysis was performed using the MaxQuant software (version 1.5.5.1) with standard settings. Used database consist of Human entries from Uniprot (reference proteome UniProt 2018_09) and 250 contaminants (MaxQuant contaminant database). Graphical representation and statistical analysis were performed using Perseus (version 1.6.1.1). A standard t-test was used to evaluate protein abundance difference between samples.

### TopBP1 expression and purification

Plasmids for protein expression in insect cells using baculoviruses are listed in Table **Plasmids**. For the production of bacmids, 50ng of pFastbac plasmids were transformed into MultiBac DH10 cells (Invitrogen, 10361-012) and positive clones were selected on LB Ampicilin (100μg/ml) plates supplemented with 40μg/ml IPTG, 100μg/ml XGal and 7μg/ml Gentamycin. Blue colonies were screened for the presence of inserts by colony PCR using pUC/M13 Forward and Reverse oligos (Bac-to-Bac® Baculoviruses Expression System_*invitrogen user guide*). To generate baculoviruses, 12×10^6^ Sf9 cells (1ml) were transfected with 5μg of purified bacmid using 15μl of Cellfectin™ (Invitrogen, P/N 58760). After 5h incubation, 9ml of medium were added to Sf9 cells and cultures were incubated for 2.5 days. The supernatant (P1) was collected by centrifugation (400 rcf 10min) and a 1/100 dilution was used to infect 4×10^6^ cells/ml Sf9 cells. Cells were incubated for 2 days and the supernatant (P2) was collected by centrifugation. Expression of fluorescent proteins was verified by Western Blotting and microscopy. For protein expression, a 1/10 dilution of freshly prepared P2 was added to 2×10^6^ cell/ml Sf9 culture and incubated for 48 to 72 hours.

Infected cells were collected and lysed mechanically using a HTU-DIGI-French-Press (10000 PSI) in 15X packed cell weight hypertonic lysis buffer (50mM Na_2_H/NaH_2_PO_4_ pH 8.0, 500mM NaCl, 1% glycerol, 0.1% CHAPS) supplemented with protease inhibitors. Lysate was clarified by centrifugation (7750 rcf, 40min, 4°C), filtered and loaded on a 5ml HisTrap HP column (GE Healthcare, 71-5027-68 AF) equilibrated with 5CV (Column Volumes) of buffer A (lysis buffer + 5mM imidazole). The column was washed with 5CV of buffer A and TopBP1^b6-8^ was eluted stepwise using 5CV of buffer A + 30mM, 50mM, 75mM, 125mM and 500mM imidazole. Peak TopBP1^b6-8^ fractions (eluted with 50mM and 75mM imidazole) were desalted (HiTrap™ or HiPrep 26/10, GE Healthcare) in physiological buffer (10mM Na_2_H/NaH_2_ PO_4_ pH 7.6, 150mM KOAc, 0.1mM MgOAc, 0.5mM DTT, 2.5% glycerol), snap frozen and stored at −80° C. Protein concentration was estimated by stainfree gel quantification using Image Lab Software.

### Nuclear extract preparation

Nuclear extracts were prepared as previously described (Vidal-Eychenie et al., 2013). HeLa S3 cells were grown to ≤ 80% confluence, collected by scrapping, centrifuged (200 rcf, 3min, 4°C) and washed twice in PBS 1X. Cell pellet was incubated on ice for 5min in 5X packed cell volume of hypotonic buffer A (10mM Hepes-KOH pH 7.9, 10mM KCl, 1.5mM MgCl_2_, 0.5mM DTT, 0.5mM PMSF) supplemented with protease (complete, EDTA free; Roche, 31075800) and phosphatase inhibitors (Fisher Scientific). Cells were then spun down (500 rcf, 5min), suspended in 2X packed cell volume of buffer A and lysed by dounce homogenization using a tight-fitting pestle. Nuclei were collected by centrifugation (4000 rcf) for 5min at 4°C, extracted in one nuclei pellet volume of buffer C (20mM Hepes-KOH pH 7.9, 600mM KCl, 1.5mM MgCl_2_, 0.2mM EDTA, 25% glycerol, 0.5mM DTT, 0.5mM PMSF) supplemented with cocktails of protease and phosphatase inhibitors, and mixed on a rotating wheel at 4°C for 30min. Nuclear extracts (supernatants) were recovered by centrifugation (16000 rcf, 15min, 4°C) and dialyzed using Slide-A-Lyzer Dialysis Cassettes (3,500-D protein molecular weight cutoff; Fisher Scientific, 68035) against buffer D (20mM Hepes-KOH pH 7.9, 100mM KCl, 0.2mM EDTA, 20% glycerol, 0.5mM DTT, and 0.5mM PMSF). Dialyzed nuclear extracts were centrifuged (100000 rcf, 30min, 4°C) to eliminate residual precipitates. The protein concentration of the clear supernatant was determined by Bradford (BioRad, C500-0205) protein assay, and aliquots were snap frozen and stored at −80°C. Important to note that the Dignam & Roeder extract preparation contain DNA fragments after centrifugation.

### TopBP1 phase separation assay

Phase separation of purified TopBP1^b6-8^ was performed in physiological buffer C (10mM Na_2_H/NaH_2_ PO_4_ pH 7.6, 150mM KOAc, 0.1mM MgOAc, 0.5mM DTT, 2.5% Glycerol). Purified TopBP1^b6-8^ WT and W1145R were digested with PreScission 3C enzyme (GenScript, Z03092-500) for 3h at 16°C to remove the 6-His and MBP tag before phase separation in reaction mixtures in buffer C containing 10μM of WT or W1145R TopBP1^b6-8^-GFP and 2% of PEG4000 (Merck-Sigma-Aldrich, 95904). Reaction mixtures were mixed by gently tapping the Eppendorf. TopBP1 droplet permeability assay was performed in two steps: first, TopBP1^b6-8^-GFP droplets were formed in reaction mixtures containing 10μM of WT TopBP1^b6-8^-GFP supplemented with 2% PEG in physiological buffer C, and then we added either 1μM of WT or W1145R TopBP1^b6-8^-RFP or *StuI/SacII* digested pX174 RFII DNA (300nM), as indicated. To study the role of DNA in the organization of TopBP1 condensates, we incubated 2.5µM of WT TopBP1^b6-8^ with 40ng 2.9 kb circular DNA plasmid DNA at 37°C during 10min in physiological buffer C. The ratio purified TopBP1:DNA was decided according to Choi JH et al., 2008. Samples were imaged on a LSM780 confocal microscope (Leica, Germany) using a 63x oil immersion objective (N.A. 1.4). DAPI and GFP fluorescence were excited at 405 and 488nm respectively, and emitted fluorescence were collected sequentially at 415-460nm and 500-550nm respectively. The pinhole size was set to 1 Airy unit.

TopBP1 phase separation in human protein extracts was performed in reaction mixtures containing 0.2μg/μl of human nuclear extract, 2μM of WT or W1145R TopBP1^b6-8^-GFP in ATR activation buffer (10mM Hepes-KOH pH 7.6, 50mM KCl, 0.1mM MgCl_2_, 1mM PMSF, 0.5mM DTT, 1mM ATP, 10μg/ml creatine kinase, 5mM phosphocreatine). Reaction mixtures were incubated for 10min at 4°C or 37°C, as indicated. 5μl of reaction mixtures were used for analyses by immunofluorescence microscopy. Fluorescence microscopy analyses were performed using PEG silanized glass slides prepared as described (Alberti et al., 2018), and coverslips were sealed with nail polish. DNA was stained with Hoechst 33258. Images were captured on an inverted microscope using a 63x objective (NA 1.4 oil).

### Immunofluorescence staining

Cells grown on coverslips were fixed with PBS/4% paraformaldehyde (PFA) for 15min at RT followed by a 10min permeabilization step in PBS/0.2% Triton X-100-PBS and blocked in PBS/3% BSA for 30min. For immunofluorescence staining, primary antibodies and appropriate secondary antibodies coupled to fluorochrome were diluted in blocking solution and incubated for 1h at RT. To detect endogenous TopBP1, cells were pre-treated with Cytoskeleton (CSK) buffer before fixation. DNA was stained with Hoechst 33258 (Invitrogen, Cat H21491) and coverslips were mounted on glass slides with Prolong Gold antifade reagent (Invitrogen, Cat P36930). Images were captured using a 63x objective (NA 1.46 oil).

### Fluorescence Recovery After Photo-bleaching (FRAP)

For FRAP experiments, *U-2-OS Tet-On* cells were seeded into μ-Dish^35 mm, high^ (Ibidi, 81156) and incubated 24h in the presence of 1μg/ml doxycycline to induce expression of eGFP-TopBP1. Imaging was realized using a 63x objective (NA 1.4). eGFP-TopBP1 bodies were photo-bleached and the GFP signal intensity of 23 spots was measured before and during 5min following photo-bleaching with an imaging frequency of 60 images/min.

### Opto-TopBP1 activation

Cells were plated at around 70% confluency in DMEM. Expression of opto-TopBP1 was induced for 16h with 2μg/ml doxycycline. For light activation, plates were transferred into a custom-made illumination box containing an array of 24 LEDs (488nm) delivering 10mW/cm^2^ (light intensity measured using a ThorLabs-PM16-121-power meter). Cry2 oligomerisation was induced using 3min of light-dark cycles (4s light followed by 10s dark). Images were captured using a 63x objective (NA 1.46 oil). A Cell Profiler (version 2.2.0) pipeline was used to quantify nuclear mCherry foci. Values were represented via violin plots or histograms elaborated in Graph Pad Prism 8. Median and quartile values were indicated by continues and dashed lines respectively. A non-parametric t-test (Mann-Withney) was used to compare mCherry spot/cell distributions between samples. For each experiment, we performed three biological replicates and we show one representative experiment.

### Live cell microscopy

Live imaging of opto-TopBP1 WT overexpressing cells was performed on an inverted microscope (AxioObserver, Carl Zeiss, Germany) using a 63x oil immersion objective (NA 1.4). Fluorescence was detected on a CMOS camera (ORCA-Flash4.0, Hamamatsu) with an exposure time set to 100ms and at frame rate of 2 images/min. All recordings were carried out at 37°C under 5% CO2. For Cry2 activation, cells were treated as explained before, under the microscope, and imaged immediately after light activation.

### Stimulated emission depletion (STED) super-resolution microscopy

For Figure 2E, *Flp-In™ T-REx™ 293* cells expressing opto-TopBP1 were exposed to blue light (488nm) to induce TopBP1 foci, as described before. For STED microscopy, immunolabelling of mCherry was realized using an anti-mCherry antibody followed by a secondary antibody coupled to Atto-647N fluorochrome. Confocal and STED imaging was performed using a quad scanning STED microscope (Expert Line, Abberior Instruments, Germany) equipped with a PlanSuperApo 100x/1.40 oil immersion objective (Olympus, Japan). Atto-647N was excited at 640nm with a dwell time of 10µs and STED was performed at 775nm. Images were collected in line accumulation mode (5 lines accumulation). Fluorescence was detected using avalanche photo diodes and spectral detection (650-750nm). The pinhole was set to 1.0 Airy units and a pixel size of 10nm was used for all acquisitions. A gating of 8ns was applied. Sphericity of opto-TopBP1 condensates was assessed using IMARIS software (Bitplane). For dual color STED imaging, Atto-647N and Abberior STAR Orange were used and respectively imaged at 640 and 561nm excitation. Detection was set to 650-750nm for Atto-647N and 570-630nm for Abberior STAR Orange. Other acquisition parameters were the same described as above.

### DNA fibers

Neo-synthesized DNA was sequentially pulse labelled with two halogenated thymidine analogs, 5-Iodo-2’-deoxyuridine (IdU at 25µM – Sigma # I7125) and 5-Chloro-2’-deoxyuridine (CldU at 50µM – Sigma # C6891), for 20min. After IdU incorporation, cells were washed two times before CldU addition. CldU incorporation (20min total) was conducted in the presence of 3min of light-dark cycles (4s light followed by 10s dark) each 7min, to assure opto-TopBP1 condensates formation and persistence. Cells were trypsinized and washed with ice cold PBS. 2000 cells were drop on top of a microscope slide and let dry at least 5min before lyse with 7µL of spreading buffer (200mM Tris.HCl pH7.5, 50mM EDTA, 0.5% SDS) during 3min. DNA was spread by tilting the slide and letting the drop running down slowly. Once air-dry, DNA spreads were fixed in methanol/acetic acid 3:1 for 10min, denatured with 2.5M HCl during 1h and blocked in PBS/1% BSA/0.1% Tween. DNA spreads were immunostained with mouse anti-BrdU, rat anti-BrdU and mouse anti-ssDNA antibodies to detect IdU, CldU and intact DNA fibers respectively. Corresponding secondary antibodies conjugated to Alexa Fluor dyes were used in a second step. Images were captured using a 40x objective (NA 1.4 oil). The acquired DNA fiber images were analyzed by using FIJI Software and statistical analysis of at least 150 IdU and CldU tracks on intact fibers was performed with GraphPad Prism 8. Mean values were indicated by red lines. One-way ANOVA analysis was applied to compare means of samples in a group.

---

**Figure S1.**
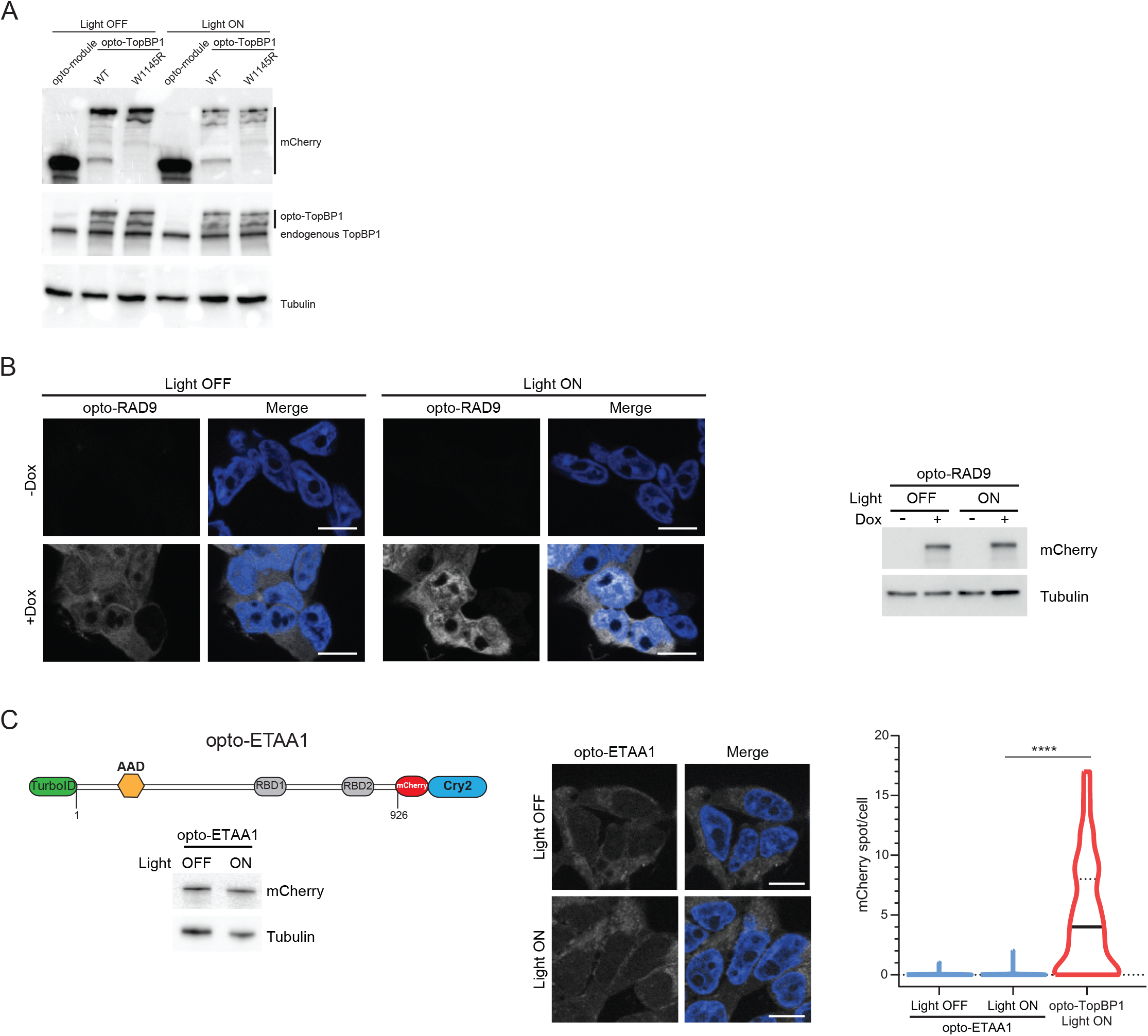
Relates to Figure 1. **A)** Cells used for optogenetic experiments in Fig 1B were analyzed by western blotting using the indicated antibodies. **B)** (Left) Representative fluorescence images of cells expressing (+Dox) or not (-Dox) opto-RAD9 construct before (Light OFF) and after (Light ON) 3min exposure to cycles of 4s light (488nm)-10s resting. (Right) Western blotting from cells used for optogenetic experiments. **C)** Schematic representation of the opto-ETAA1 with representative fluorescence images of cells expressing the respective optogenetic construct before (Light OFF) and after (Light ON) 3min exposure to cycles of 4s light (488nm)-10s resting. Western blotting of the indicated proteins is represented. Violin plot represents the number of mCherry foci per cell. Median and quartile values are represented by continues and dashed lines respectively. The statistical significance of the difference in mCherry spot/cell distributions between samples is represented by *. **B**-**C)** DNA stained with Hoechst 33258. Scale bars: 10μm.

**Figure S2.**
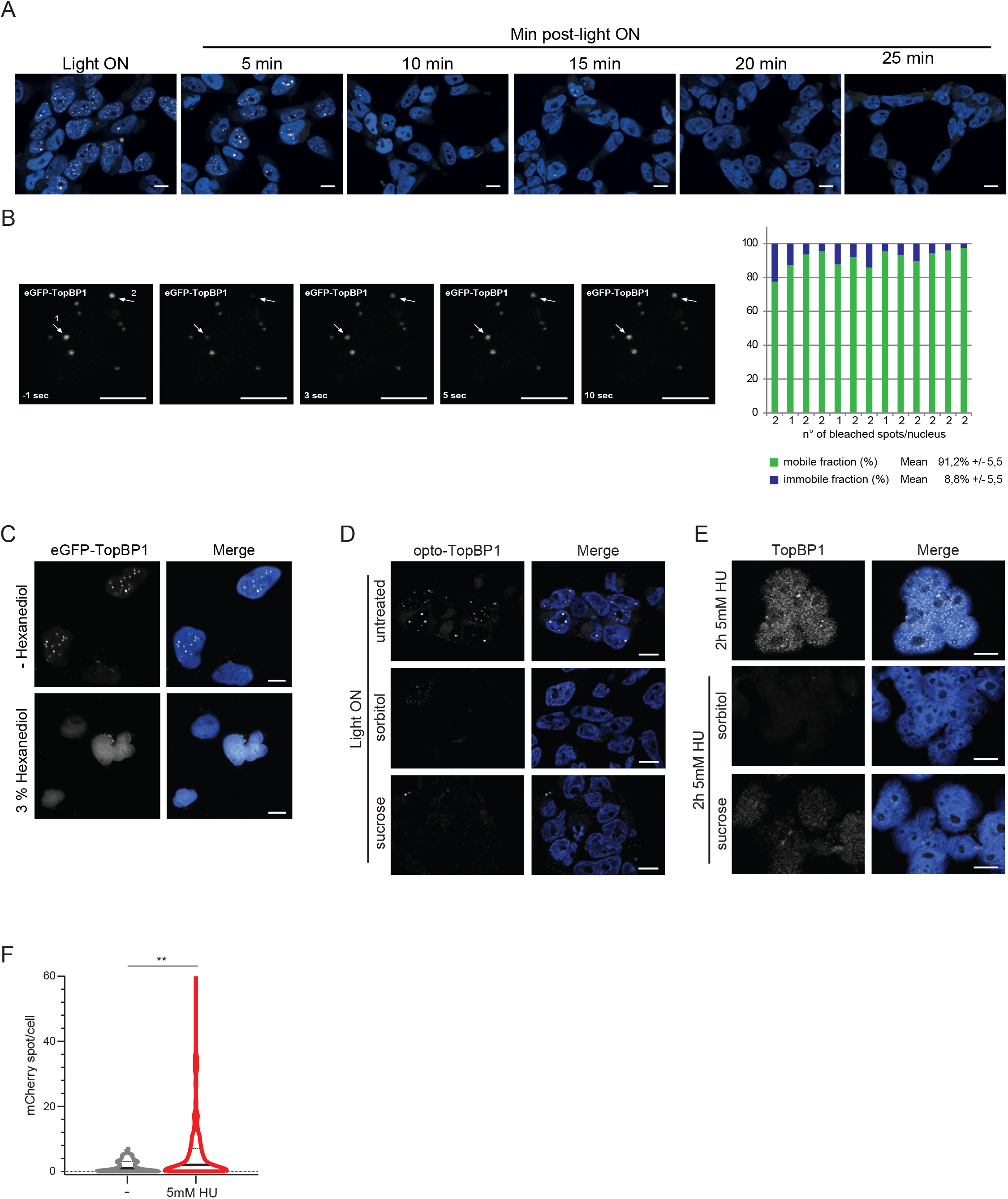
Relates to Figure 2. **A)** Representative fluorescence images of cells used for experiments in Figure 2C. DNA was stained with Hoechst 33258. Scale bars: 10μm. **B)** (Left) Fluorescence Recovery After Photobleaching (FRAP) analyses of eGFP-TopBP1 bodies expressed in *U-2-OS* cells. FRAP recordings of two individual eGFP-TopBP1 bodies are reported as examples. Scale bars: 10μm. (Right) Histogram representation of the mean of immobile and mobile fractions per nucleus (13 nuclei, 23 individual spots). **C)** Representative fluorescence images of eGFP-TopBP1 expressing *U-2-OS* cells treated, when indicated, with 3% of 1,6 Hexanediol for 5min. **D-E**) Representative fluorescence images of cells used for experiments in Figure 2D (**D**) and 2E (**E**). DNA was stained with Hoechst 33258. Scale bars: 10μm. **F)** Violin plot representing the number of mCherry foci per cell in cells expressing opto-TopBP1 WT treated with 5mM HU for 2h and not exposed to blue light. TopBP1 foci were identified with anti-mCherry antibody after CSK treatment. Median and quartile values are represented by continues and dashed lines respectively. The statistical significance of the difference in mCherry foci/cell distributions between samples is represented by *.

**Figure S3.**
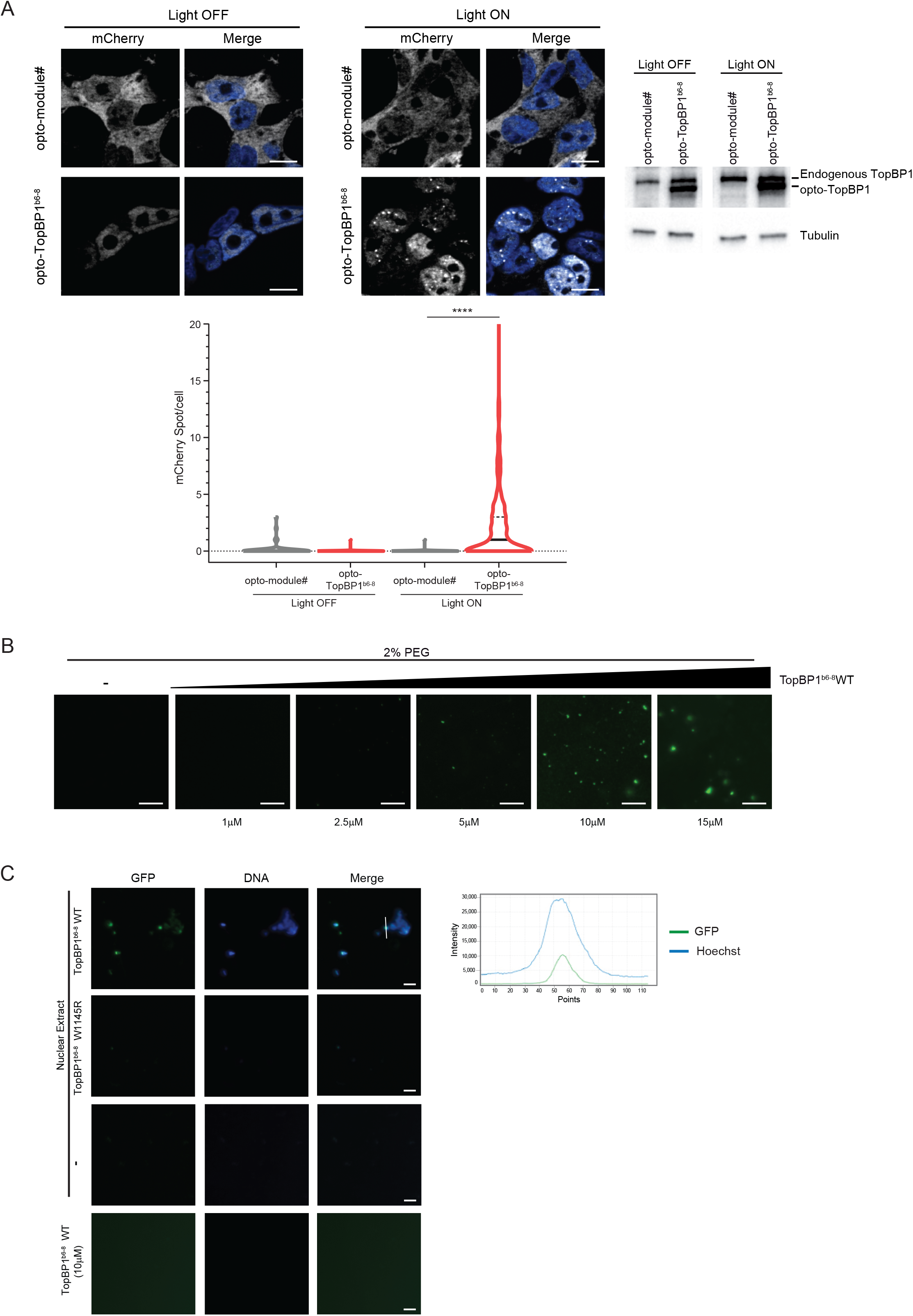
Relates to Figure 3. **A)** Representative fluorescence images of cells expressing opto-TopBP1^b6-8^ WT before (Light OFF) and after (Light ON) 3min exposure to cycles of 4s light (488nm)-10s resting. Control cells express the opto-module# mCherry-Cry2. DNA was stained with Hoechst 33258. Western blotting of the indicated proteins is represented. Violin plot represents the number of mCherry foci per cell. Median and quartile values are represented by continues and dashed lines respectively. The statistical significance of the difference in mCherry spot/cell distributions between samples is represented by *. (**B**) Representative fluorescence microscopy images obtained with increasing concentrations of TopBP1 in the presence of phosphate buffer pH 7.6 containing 150mM KAc and 2% PEG. (**C**) Representative fluorescent images of Figure 3F with DNA and merge panels. Images of condensates in suspension are represented. DNA was stained with Hoechst 33258. Line scan of GFP-Hoechst signals is used to analyze co-localisation. (**A-B-C**) Scale bars: 10μm.

**Figure S4.**
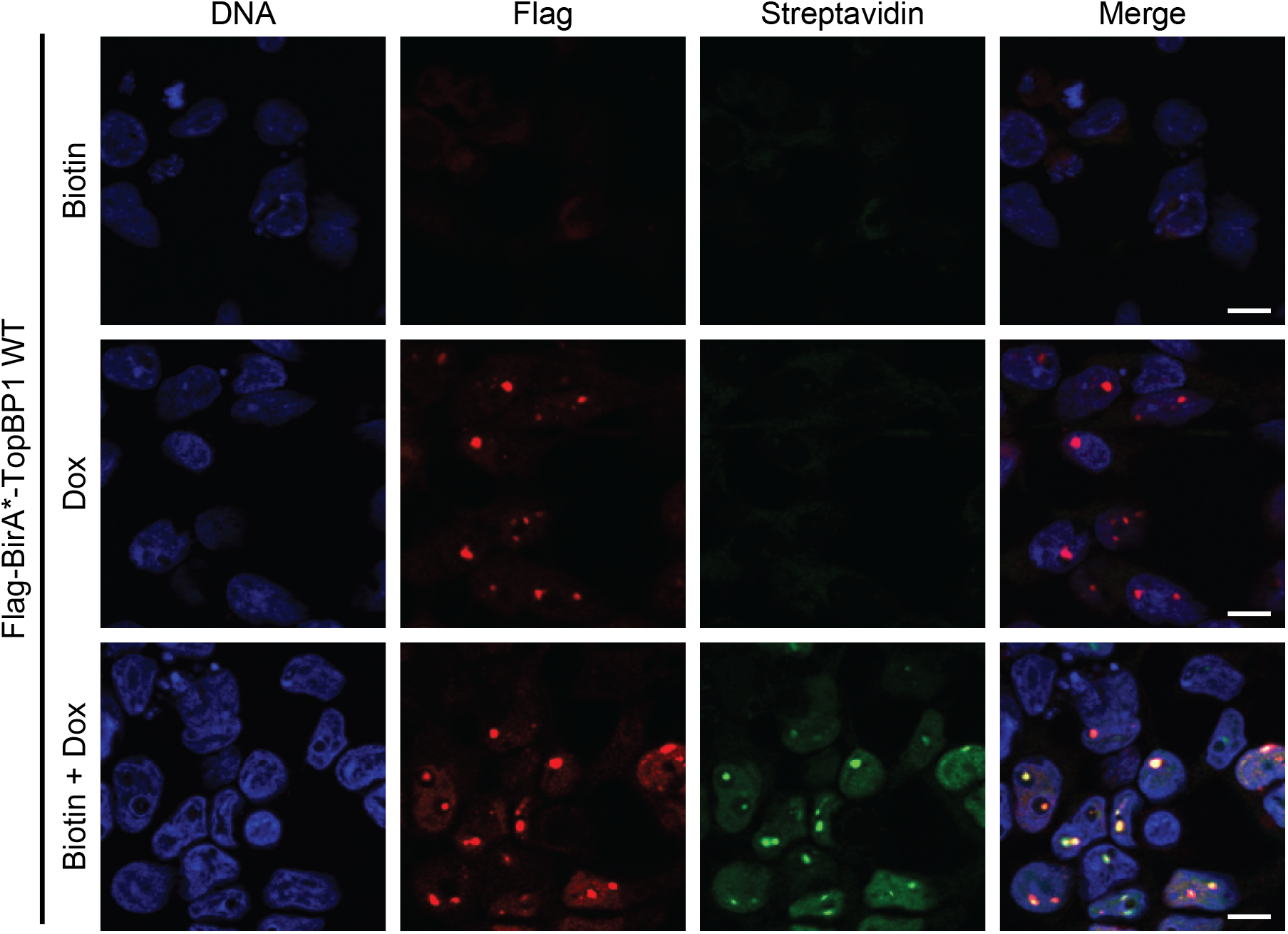
Relates to Figure 4. Immunofluorescence staining of Flag-BirA*-TopBP1 WT using anti-Flag antibody. Biotin conjugates were revealed using AlexaFluor streptavidin. When indicated, cells were grown in the presence of 1µg/ml doxycycline and 50µM biotin. DNA was stained with Hoechst 33258. Scale bars: 10μm.

**Figure S5.**
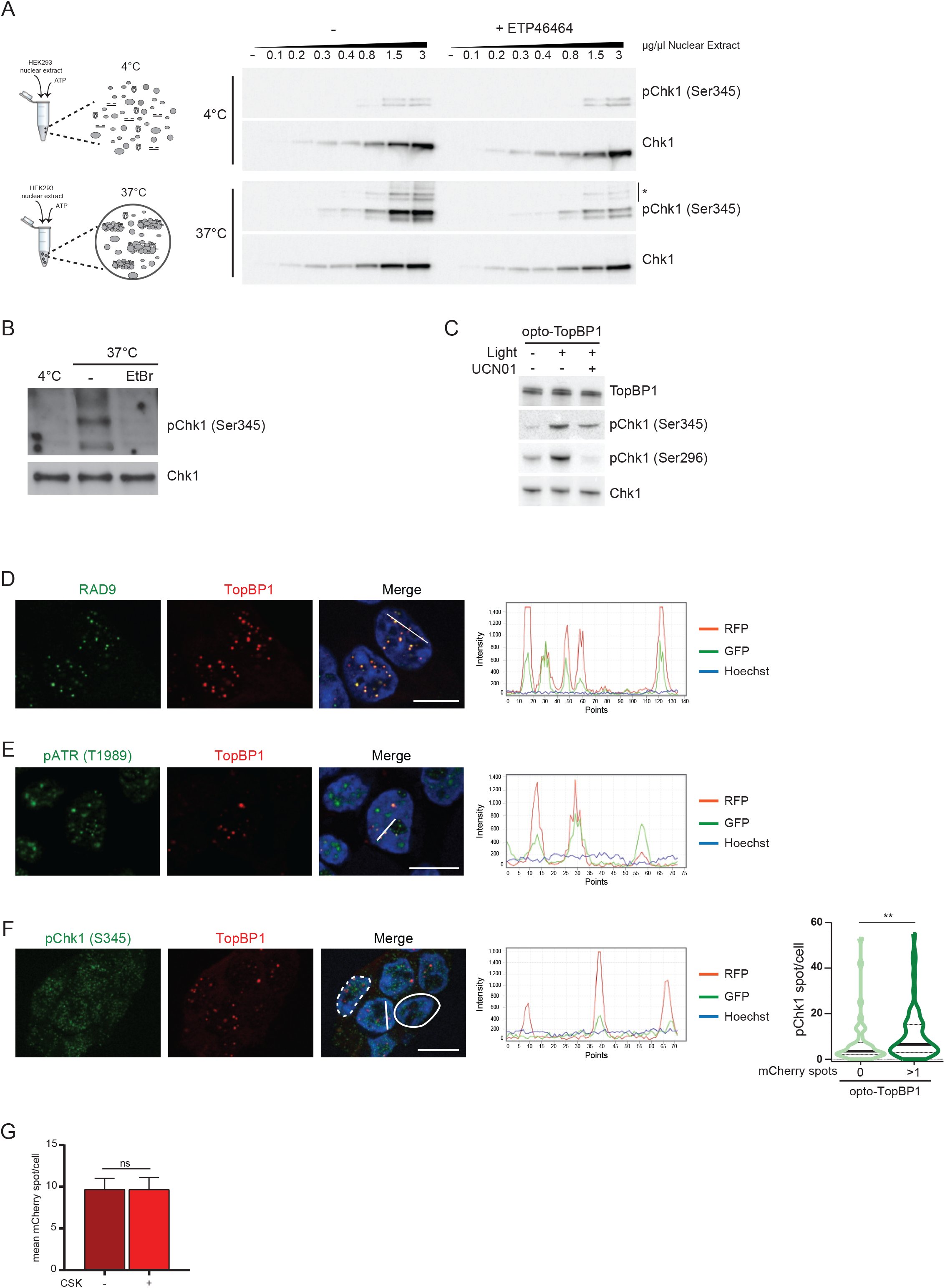
Relates to Figure 5. **A)** Western blot analysis of Chk1/phospho Chk1 (Ser345) in reaction mixtures containing increasing concentration of “Dignam and Roeder” nuclear extracts, and, when indicated, the ATR inhibitor ETP-46464 (10μM). Reaction mixtures were incubated for 10min at 4°C or 37°C, as indicated, in the presence of 150mM KAc in phosphate buffer pH 7.6. * represents unspecific bands. The design of the experiment is represented schematically. **B)** Western blot analysis of Chk1/pChk1 (Ser345) in reaction mixtures containing 4µg/µl of “Dignam and Roeder” nuclear extracts in phosphate buffer pH 7.6 supplemented with 150mM KAc. Reaction mixtures were incubated for 10min at 4°C or 37°C and pre-incubation of extracts with ethidium bromide (EtBr) is indicated. **C)** Western blot analysis of TopBP1 and Chk1/pChk1 (Ser345-Ser296) of cells expressing opto-TopBP1, exposed to 3 minutes cycles of 4s light (488nm)-10s resting. When indicated, cells were pre-treated with 10μM Chk1 inhibitor UCN-01. **D-E-F)** Immunofluorescence staining of opto-TopBP1 expressing cells activated by light and RAD9 (**D**) pATR (Thr1989) (**E**) and pChk1 (Ser345) (**F**) using specific antibodies. Line scans are used to analyze co-localisation. DNA was stained with Hoechst 33258. Scale bars: 10μm. Violin plot (**F-**right) representing spot of pChk1 in cells with (dashed white line) or without (solid white line) opto-TopBP1 foci. **G)** Histograms representing the mean of mCherry foci per cell. Cells expressing opto-TopBP1 WT were first exposed for 3min to cycles of 4s light (488nm)-10s resting (Light ON) and then treated (+) or not (-) with Cytoskeleton (CSK) buffer to keep chromatin-bound proteins.

**Figure S6.**
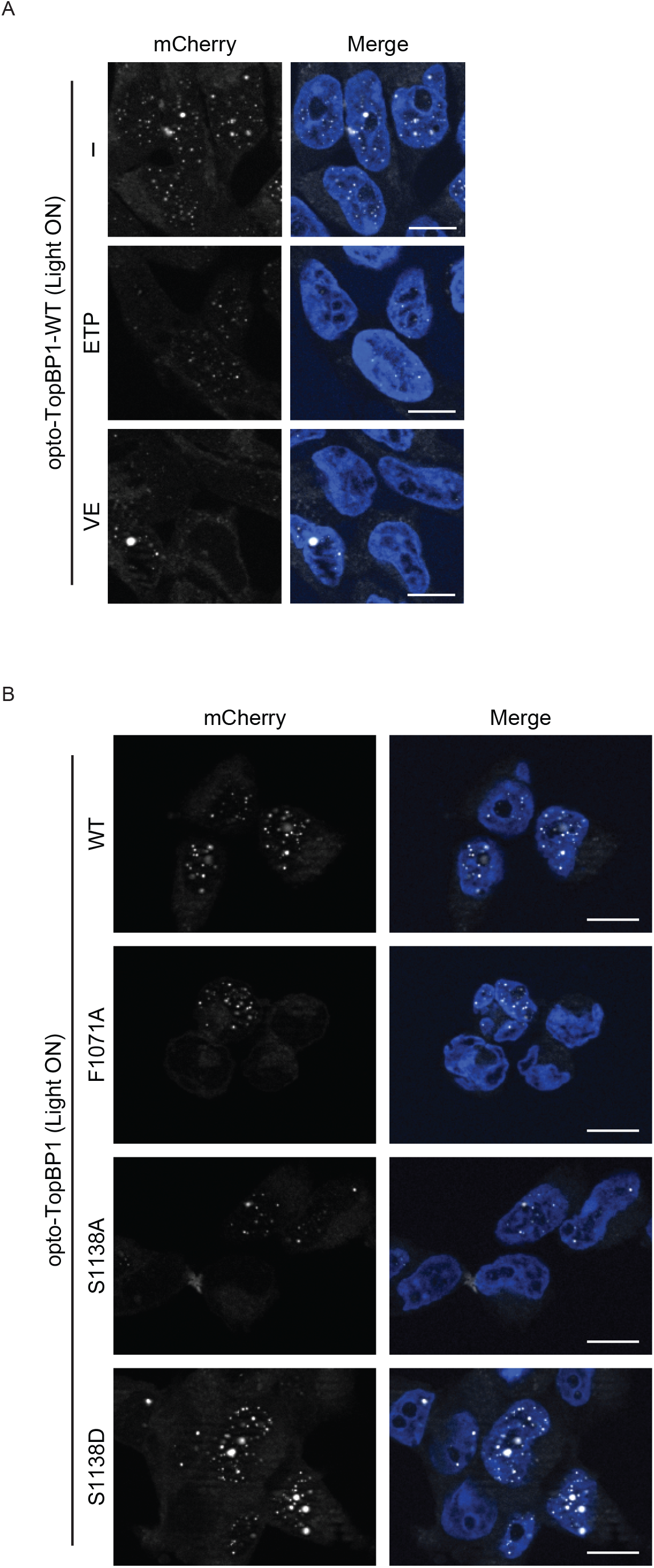
Relates to Figure 6. **A**) Representative fluorescence images of cells used for optogenetic experiment in Figure 6A. **B**) Representative fluorescence images of cells used for optogenetic experiment in Figure 6C.

